# Molecular architecture and diversity of StopGo/2A translational recoding

**DOI:** 10.1101/2025.11.07.687186

**Authors:** Xueyan Li, Philipp K. Zuber, Gary Loughran, Pramod R. Bhatt, Fatema Alquraish, V. Ramakrishnan, Andrew E. Firth, John F. Atkins

## Abstract

Viral 2A sequences trigger a co-translational peptide bond formation “skipping” event, termed “StopGo”, to generate two separate proteins from a single open reading frame without classical termination. To investigate the mechanism of StopGo, we determined the cryo-EM structure of a mammalian ribosome positioned at the foot-and-mouth disease virus 2A (F2A) site. The structure shows how interactions between the F2A nascent chain (NC) and the ribosomal exit tunnel induce a conformational change in the peptidyl transferase center that precludes further translation elongation but instead pre-exposes the P-tRNA:F2A-NC ester bond for hydrolysis and NC release. Additionally, we bioinformatically characterized variation and host association across nearly 10,000 StopGo sequences identified in virus genomes. We expanded the canonical core motif to (D/G/C/N)(V/I)ExNPGP and identified additional rare but functional variants. We also revealed several distinct upstream motifs that we showed biochemically to be important for StopGo activity. Interestingly, although StopGo is known to be functionally active in plants, we found no evidence for natural utilization of StopGo by plant viruses. Overall, these findings provide valuable insights into a unique translation recoding mechanism, and lay foundations for further optimization of multi-gene expression in biotechnology.

## Introduction

Translational recoding allows ribosomes to reinterpret mRNA sequences in ways that expand the coding potential of genomes (1-3). A particularly intriguing example involves “2A” sequences, which promote (i) protein release without termination on a stop codon, and (ii) continuation of translation without re-initiation to synthesize a separate downstream encoded protein (4-7). The phenomenon was originally discovered by M. Ryan and colleagues to occur following synthesis of the C-terminal end of the foot-and-mouth disease virus (FMDV) 18 amino acid (aa) 2A protein. The responsible sequence is therefore often referred to as “F2A” (8-10). Though initially termed “2A”, the phenomenon was later found in other viral sequences not homologous to the FMDV 2A protein. Here, we use the designation “StopGo” to reflect both (i) the mechanism of action and (ii) the natural variety of corresponding sequences ranging well beyond just picornavirus 2A peptides (6, 11).

Natural StopGo utilization is largely confined to viral decoding (12) and, as its enabling motifs only very rarely occur in cellular coding sequences (absent in mammals) (13), it may constitute an anti-viral target (14). Early work showed that, in addition to FMDV, other picornaviruses such as porcine teschovirus (“P2A”), equine rhinitis A virus (“E2A”) and the cardioviruses encephalomyocarditis virus (EMCV) and Theiler’s murine encephalomyelitis virus (TMEV) utilize StopGo sequences in a central region of their polyprotein between upstream capsid and downstream replication protein sequences (10, 15, 16). Many other RNA viruses utilize StopGo to process other polyproteins (17, 18). Whereas StopGo sequences function in all tested eukaryotic systems, they do not function in *E. coli* (in *E. coli*, the closest phenomenological counterpart is non-programmed and mechanistically distinct (19)). Importantly, StopGo sequences retain their functionality when grafted between any two proteins (20, 21). This has made this non-proteolytic mechanism of polycistronic protein expression from just one continuous ORF a valuable tool for research and biotechnology (22-24), including cases of co-expression of all the proteins of a pathway (25, 26), and trimming test sequences off reporter proteins to prevent assay artifacts (27).

At its core, StopGo features Pro-Gly↓Pro codons in a specific sequence context. The peptide bond formation “skipping” event (indicated by ↓) separates the polyprotein between Gly and the second Pro. The PG↓P amino acids form the C-terminal end of a highly conserved octapeptide motif, -D(V/I)ExNPG↓P that is critical for StopGo activity (4). Sequences encoded further upstream also have an important influence on efficiency (28, 29). With sufficient upstream native sequence, the amount of unseparated product is extremely low for both EMCV and FMDV (10, 12, 29, 30). StopGo activity is not conferred through specific mRNA structures but relies on interactions between the StopGo nascent chain (NC) and the ribosomal peptidyl transferase center (PTC) (7, 28). However, direct evidence of the interactions and re-arrangements of the PTC, and a detailed mechanistic characterization of the StopGo phenomenon is lacking. Furthermore, the extent of natural variability within the core octamer, besides further upstream sequences, has not been studied in the context of the enormous amount of new virus sequencing data generated recently by metagenomic and other studies.

Here, we integrate structural biology, functional assays and bioinformatic analysis to investigate the mechanism and sequence diversity of StopGo. We report the cryo-EM structure of a mammalian ribosome stalled at the F2A sequence, revealing that interactions between the F2A- NC and the ribosomal peptide exit tunnel play a dual role in translation regulation through (i) contributing to ribosomal pausing by stabilizing the NC in the exit channel, potentially delaying elongation at a critical moment, and (ii) inducing structural rearrangements in the PTC that prevent addition of the next amino acid while pre-exposing the tRNA:F2A ester bond, facilitating its hydrolysis and subsequent NC release. A comprehensive bioinformatic analysis of nearly 10,000 viral StopGo sequences identifies rare variants within the core octapeptide, and several distinct functional upstream motifs, both of which we experimentally verify.

## Results

### Cryo-EM structure of the StopGo/F2A peptide bound translational complex

To understand the mechanism by which StopGo sequences induce translational peptide bond “skipping”, we sought to obtain structural insights into the translation complex positioned at the F2A site by single particle cryo electron microscopy (cryo-EM). Since StopGo is too rapid to enable isolation of the transient WT target complex for structural investigation (7), we substituted P19 of the F2A motif with a stop codon, creating an F2A PGX mRNA (Fig. 1A). The construct encodes an N-terminal 3X-FLAG tag linked to the native FMDV genomic sequence comprising the C-terminal 32 aa of virus protein 1 (VP1) followed by the 18 aa of protein 2A and the UGA stop codon (X). Translating the mRNA in vitro in presence of the dominant negative AAQ mutant of eukaryotic release factor 1 (eRF1) then allows site-specific stalling of the translating ribosome at the StopGo site. Notably, positioning and conformation of the P-site tRNA, NC and peptidyl transferase center (PTC) in structures of elongation complexes, or termination complexes prior to accommodation of eRF1 are identical to complexes containing eRF1-AAQ accommodated in the A-site (31). This indicates that the presence of eRF1-AAQ (and its mutated catalytic motif) has no effect on the P-tRNA:NC and the resulting complex therefore accurately reflects the pre-release state of the StopGo complex.

**Figure 1.**
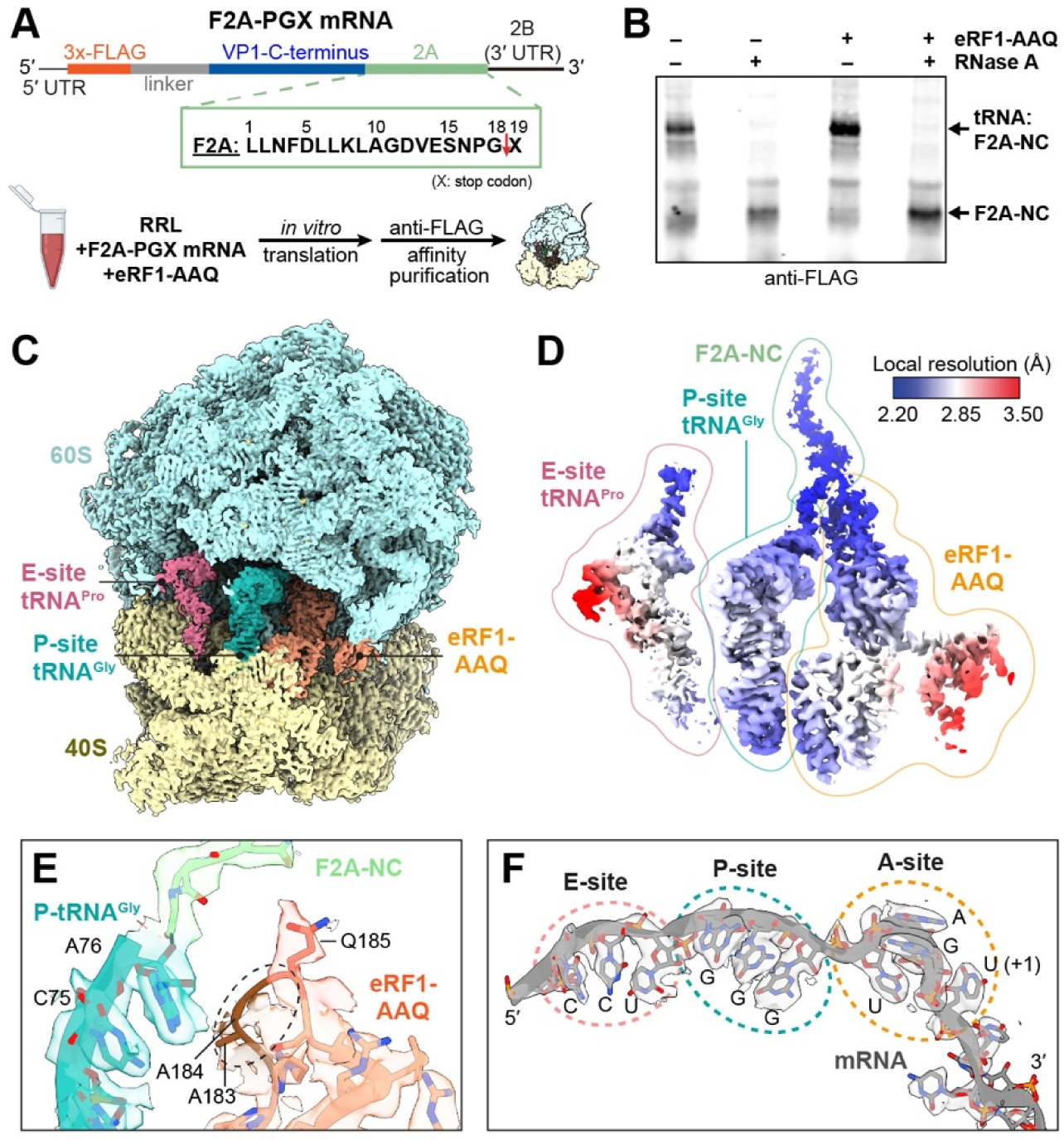
Cryo-EM structure of the F2A-StopGo complex. **(A)** Schematic of the F2A-PGX mRNA (top; X = stop codon) and complex preparation strategy (bottom). **(B)** Anti-FLAG Western blot of in vitro translation reactions of the F2A-PGX mRNA in absence/presence of eRF1-AAQ. The free and tRNA-bound F2A-NC species are labelled. **(C), (D)** CryoEM maps of the complete F2A-StopGo complex (C), or E- and P-site tRNAs, F2A-NC and eRF1-AAQ, coloured based on local resolution (D). **(E), (F)** Map and model of P-site tRNA^Gly^, the attached F2A-NC, and eRF1- AAQ in the PTC region (E) or the mRNA (F). The disordered region around the eRF1-AAQ motif (E) and the A, P and E-sites (F) are highlighted.

Previous studies had shown that the F2A-P19X mutation itself slows release of the F2A-NC from the tRNA, suggesting NC-induced structural changes that impair regular translation termination (28). Consistently, in vitro translation of the F2A-PGX mRNA in unmodified rabbit reticulocyte lysate (RRL) showed accumulation of the ribosome stalled at the StopGo site, as indicated by a tRNA:F2A-NC band in an anti-FLAG Western blot (Fig. 1B). Addition of eRF1-AAQ further increased the amount of stalled ribosomes (Fig. 1B) and stabilized the complex during FLAG affinity purification. Subsequent cryoEM analysis then allowed us to resolve the F2A-NC bound state after focused classification on eRF1-AAQ and the P-site tRNA:NC at an overall resolution of 2.6 Å (Fig. 1C, Fig. S1, Table S1).

The cryo-EM density reveals the F2A-NC linked to tRNA^Gly^ in the ribosomal P-site (Fig. 1D, 2A), while the E-site tRNA^Pro^ is more loosely engaged (Fig. 1D). The mRNA is clearly resolved and allows unambiguous identification of the ribosome at the StopGo site (Fig. 1F). eRF1-AAQ is accommodated in the A-site, bound to the UGA stop codon and the following 3′ mRNA nucleotide as observed previously (31). As expected, the Ala residues of its AAQ motif are largely disordered and do not contact the tRNA^Gly^:F2A-NC ester bond (Fig. 1E).

### Interactions of StopGo/F2A peptide with the ribosomal exit channel

The cryo-EM map clearly resolves the tRNA^Gly^ in the P-site and the F2A-NC within the ribosomal peptide exit tunnel, providing a detailed snapshot of the stalled complex. Notably, F2A residues L6–G18 can be fitted well within the NC density; the preceding residues are more flexible and hence less well defined (Fig. 2A). The tRNA^Gly^-CCA:G18 ester bond is clearly formed (Fig. 2A), indicating that the complex is captured just before NC release, a critical step in StopGo.

**Figure 2.**
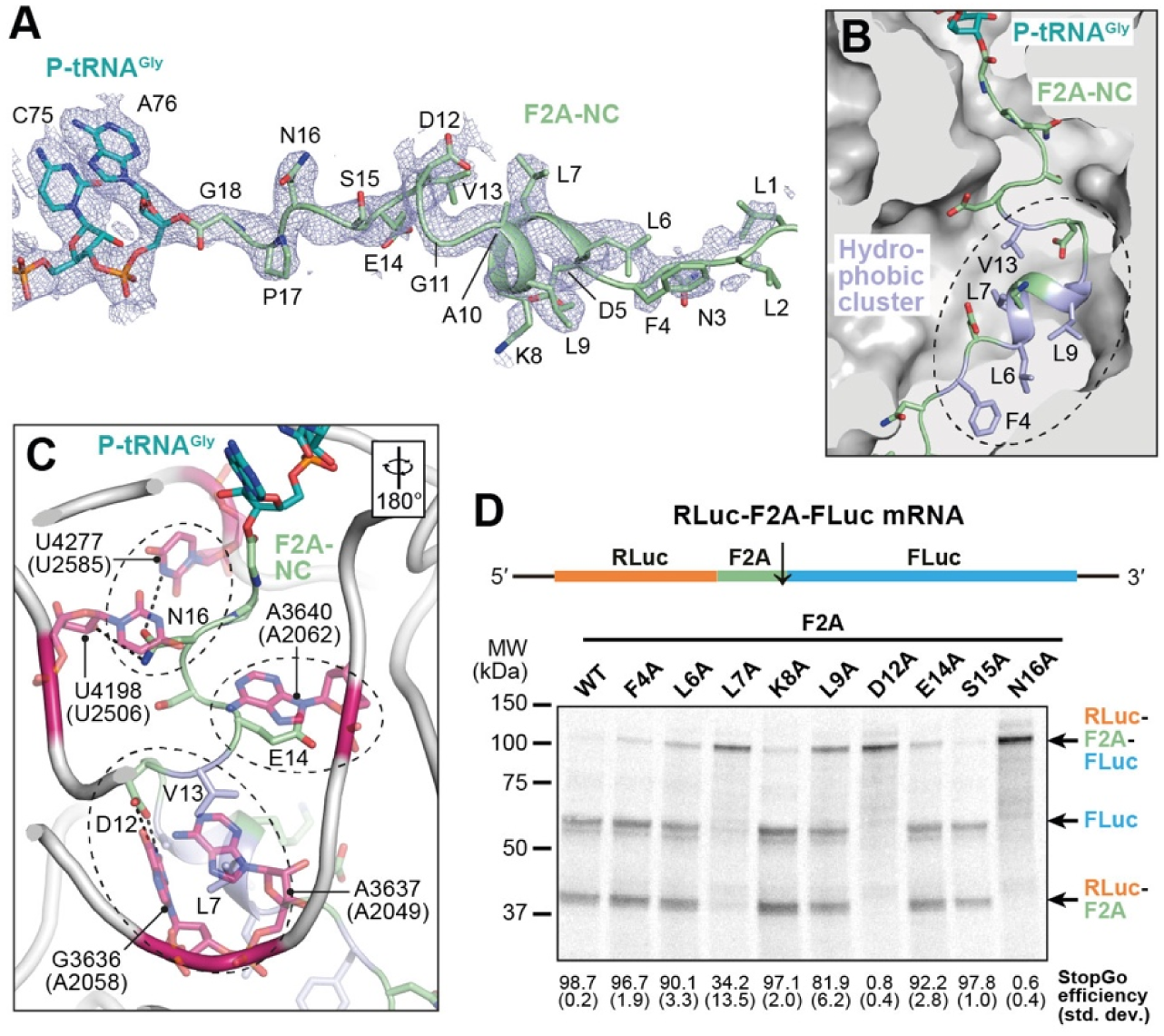
Interactions of the F2A-NC and the ribosomal peptide exit tunnel. **(A)** Cryo-EM density and model of tRNA^Gly^-CCA end and attached F2A-NC. Boxed: enlargement of the region around the helical segment of the F2A-NC. **(B)** Path of the F2A-NC in the ribosomal peptide exit tunnel. Residues constituting a hydrophobic cluster are highlighted **(C)** Polar interactions between F2A-NC side chains and 28S rRNA residues. Homologous rRNA nucleotides of *E. coli* are given in parentheses; the orientation relative to (B) is indicated. **(D)** Autoradiograph of SDS-PAGE gel showing StopGo activity assay of F2A single mutations using a dual reporter construct (top). The calculated StopGo efficiency (%) and standard deviations (±%) are given below (*n* = 3).

The defined conformation of the F2A-NC is stabilized through several specific contacts with 28S rRNA residues in the nascent peptide exit tunnel (Fig. 2B, C). At the tunnel entrance, N16 establishes hydrogen bonds with N3 of U4277 and the 2′ OH group of U4198 in the 28S rRNA. Further into the tunnel, the side chain of E14 forms a stacking interaction with the base of A3640, positioning it near the ribosomal RNA for potential stabilization of the local architecture. The side chain of D12 forms hydrogen bonds with the Watson-Crick edge of G3636 in the 28S rRNA. (Fig. 2C). Additionally, the FDLLKL motif in the middle of the F2A-NC adopts a helical conformation that establishes a hydrophobic cluster (Fig. 2A), contributing additional Van-der-Waals interactions with the exit tunnel (Fig. 2B). In particular, L7 is wedged between the bases of G3636 and A3637, aiding the stabilization provided by D12 (Fig. 2C). Altogether, these interactions clamp the F2A-NC in a specific conformation within the ribosome’s nascent peptide exit tunnel.

To assess the functional relevance of the interactions between F2A-NC and exit tunnel, we performed StopGo activity assays using a reporter construct that contains a single ORF encoding an upstream Renilla luciferase (RLuc) and downstream Firefly luciferase (FLuc) separated by the F2A sequence. In vitro translation in RRL using ^35^S-Met labelling followed by SDS-PAGE and autoradiography showed that the WT F2A sequence exhibited a StopGo efficiency of >98% (Fig. 2D). Alanine substitution of F2A N16 completely abolished StopGo activity, highlighting its essential role for StopGo skipping. Similarly, D12A abolished the activity. Additionally, single mutations of the hydrophobic residues (F4A, L6A, L7A, K8A, L9A) affected activity to varying degrees. L7A mutation reduced activity to 34%, demonstrating its critical role in exit channel engagement, while L9A mutation reduced activity to 82%, indicating a significant but less critical contribution. These findings are broadly in agreement with previous alanine scanning experiments (28) using different upstream and downstream sequences, and confirm that both hydrogen bonding and hydrophobic interactions within the ribosomal exit channel are crucial for F2A-mediated StopGo.

### Inactivation of the PTC in the F2A-StopGo complex

To investigate the functional state of the PTC in the F2A bound translational complex, we compared our structure to a termination complex in the pre-release state containing accommodated eRF1-AAQ (PDB: 5LZV) (31). Superposition of the two models shows conformational differences at the tRNA^Gly^ CCA end, the path of the NC, and at the PTC (Fig. 3A-C). In our F2A StopGo complex, the CCA end of the P-site tRNA^Gly^ maintains normal base pairing with the P-loop. However, the CCA sugar-phosphate backbone, and with it the remainder of the tRNA, is shifted out towards the solvent by about 3–4 Å compared to the canonical termination complex. This repositioning is likely caused by the conformation of the NPG residues at the C-terminus of F2A. In canonical complexes, the NC C-terminus adopts an extended beta-strand conformation, whereas in the F2A-NC, G18 and P17 adopt an irregular loop conformation. This forces the NC further into the catalytic cleft of the PTC, below the trajectory observed in canonical complexes. Furthermore, the altered geometry displaces the carbonyl group of the ester bond – normally targeted by the α-amine of the incoming aminoacyl-tRNA – away from the catalytically active position. Modelling a tRNA^Pro^ in lieu of eRF1-AAQ in the A-site shows a carbonyl:α-amine distance of ∼6.9 Å, that strongly contrasts with the 3.2 Å spacing observed in regular pre- transpeptidation complexes (Fig. 3D) (32). This large separation between the two reactive groups effectively prevents addition of the next amino acid. Overall, the geometry of the F2A-NC thus levers the tRNA^Gly^:G18 ester bond out of its normal, catalytically competent position within the PTC. We note that, since the structural differences can be deduced from comparison to other eRF1-AAQ stalled complexes, and since no contacts between eRF1-AAQ and the tRNA or NC were observed, the functional state of the ribosome in our F2A-StopGo complex is not a result of eRF1-AAQ binding but is instead solely induced by the F2A-NC itself.

**Figure 3.**
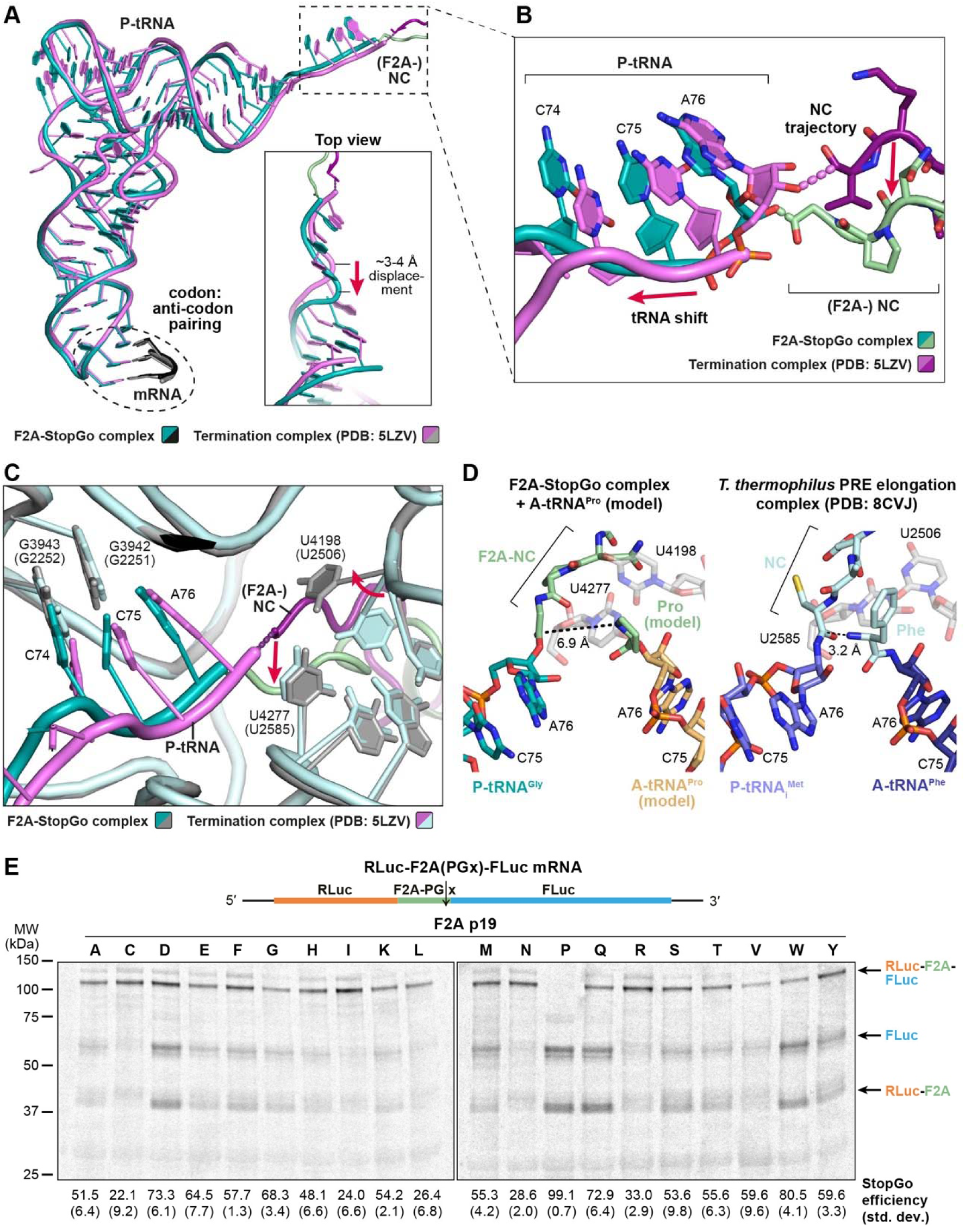
Inactivation of the PTC in the F2A-StopGo complex. **(A)** – **(C)** Superposition of the models of the F2A-StopGo complex and a canonical termination complex in the pre-release state, trapped with eRF1-AAQ (PDB: 5LZV), showing the P-tRNA/NC (A), close-up view of the CCA end/NC C-terminus (B), or PTC (C). Models are colored as indicated; structural changes to the positioning of the P-tRNA, trajectory of the NC or rRNA residues of the PTC imposed by the F2A-NC are indicated by arrows. *E. coli* numbering of rRNA residues is given in parentheses. **(D)** Positioning of the P- and (modelled) A-site tRNAs within the PTC of the F2A-StopGo complex (left) or a bacterial PRE elongation complex in the pre-transpeptidation state (right, PDB: 8CVJ). The A-tRNA^Pro^ was modelled based on the A-site tRNA of PDB model 6Y0G. The distances between the ester bond and amino group of the A-site aminoacyl-tRNAs are indicated. **(E)** Autoradiograph showing StopGo activity assay of 20 amino acid substitutions of F2A P19 using a dual reporter construct (top). The calculated StopGo efficiency (%) and standard deviations (±%) are given below (*n* = 3).

In addition to the shift of the P-tRNA^Gly^ CCA end and ester bond, the conformation adopted by the F2A-NC permits N16 to engage in hydrogen bonding with 28S rRNA residue U4277 (U2585) and a stacking interaction with U4198 (U2506; residues in parentheses indicate *E. coli* numbering) (Fig. 2C). Structural studies of peptide bond formation in prokaryotes have shown that the two homologous residues are at the PTC-end of a conformational change that senses tRNA binding to the A-loop (33). In the absence of accommodated A-tRNA, the PTC adopts an “uninduced” state, in which U2585 laterally covers the tRNA:NC ester bond, thus protecting it from spontaneous hydrolysis (Fig. S2A, B). Upon accommodation of a charged tRNA, U2585 slides away, allowing correct positioning of the ester bond and α-amine of the A-site tRNA for nucleophilic attack (“induced” state (33)). The side chain of U2506 is more flexible and can sample multiple conformations. In some structures of the uninduced state, it was observed to base-pair with G2583 (Fig. S2A), whereas in others it interacts with U2585 (Fig. S2B) (34). However, as a general observation, any interactions are broken and U2585 shifts to enable correct positioning of the A-site aa in the induced state. In structures of uninduced mammalian elongation/termination complexes, U4277 and U4198 loosely interact with each other (Fig. S2C); tRNA accommodation leads to the same shift for U4277 that uncovers the ester bond, and perturbation of interactions with U4198 (Fig. S2D; note that the only available high-resolution structure of a mammalian PRE state (PDB: 6Y0G) likely reflects the product state, with NC attached to A-tRNA). Interestingly, in the F2A-StopGo complex, the interaction of N16 forces U4198 into a specific conformation, that is turned about 90° relative to the uninduced state, but also distinct from the conformation in the induced state structure (Fig. S2E). Comparison with the pre-release termination complex further shows that swapping of U4198 is likely triggered by a clash with F2A-N16 (Fig. S2F), and then stabilized in the resulting conformation through stacking with N16 and the additional H-bond of N16 with U4277. The F2A-NC thus remodels the PTC into a distinct state that shifts U4198 into a conformation that is likely unable to support correct positioning of the incoming acylated tRNA for nucleophilic attack on the ester bond.

### Role of the incoming A-site tRNA in StopGo/F2A-NC release

Our strategy to trap the F2A complex relies on substitution of the P19 codon, which would normally reside in the A-site, by a stop codon to recruit dominant-negative release factor eRF1- AAQ. It is important to emphasize that eRF1 was shown to be neither required for StopGo activity (since an in vitro system lacking eRF1 and eRF3 is perfectly able to support ribosomal “skipping”) nor able to release the NC, but instead prevents hydrolysis (Fig. 1B) (28, 35). This suggests that the incoming tRNA^Pro^ plays a pivotal role in the NC release reaction.

We therefore sought to assess the importance of the identity of the incoming A-tRNA by performing StopGo activity assays using the F2A dual reporter construct with 19 different aa substitutions at p19 (Fig. 3E). As expected, Pro at p19 exhibited the highest activity. However, substitutions with Asp, Gln or Trp retained over 70% activity, and more than half of all substitutions maintained over 40% activity. Thus, these data suggest that while Pro is optimal, the identity of the incoming tRNA is not strictly deterministic for StopGo activity. The sidechain of the A-tRNA aminoacyl moiety therefore likely plays a passive role in peptide release. For instance, the higher efficiency of peptide release mediated by Pro is likely due to its rigid, cyclic sidechain, which makes its amino group a generally poor nucleophile for transpeptidation (36). This low reactivity prolongs the residence of tRNA^Pro^ in the A site and may provide a kinetic window for rearrangement of the StopGo-NC/PTC into their distinct inactive state, and for subsequent NC- hydrolysis to occur. In addition to playing a passive role for hydrolysis, the A-tRNA could act as a positional cofactor that actively promotes NC-release, e.g. by coordinating a water molecule (see Discussion). However, further mechanistic studies will be needed to dissect how the A-tRNA and its identity modulate this finely balanced reaction.

### Prediction of permissible variants of the StopGo core octapeptide

The canonical StopGo octameric core motif has been defined as D(V/I)ExNPGP, where x represents any amino acid (7, 28). Prior studies have also noted a preference for a G preceding D12 as well as confirmed functional D12G and D12C variants (12). To systematically explore potential naturally occurring sequence variations of the core StopGo motif, we searched proteomes for octapeptides that either matched D(V/I)ExNPGP exactly, or differed at a single position. We reasoned that, if the observed number of occurrences of any such octapeptide was much greater than the expected number of occurrences among random sequences, then the observed octapeptide would likely represent a functional variant of StopGo (albeit possibly with altered activity). This approach would be more efficient in proteomes with higher frequencies of StopGo utilisation as this would lead to larger differences between observed and expected counts. Thus, since StopGo appears to be used commonly in RNA virus gene expression but only rarely by cellular organisms (13, 17) we turned to RNA virus proteomes.

We compiled 869,685 non-redundant viral nucleotide sequences from NCBI and additional datasets (37-41). After extracting open reading frames ≥1500 nt, we obtained 793,449 theoretical aa sequences. Given the overrepresentation of certain viral taxa in the database (e.g., FMDV), we clustered sequences at 70% identity (CD-HIT), yielding 131,941 representative sequences. From these, we identified 908 exact matches to D(V/I)ExNPGP and 1557 octapeptides differing at a single position. Most (83%) of the inexact matches corresponded to the previously known variants mentioned above, i.e. p12 = G (1207 cases) and p12 = C (85 cases).

Analysis of the 2465 octapeptides (Fig. 4A) confirmed that p14, p16, p17, p18 and p19 are almost exclusively E, N, P, G and P, respectively, whereas V and I are both common at p13, and p12 is predominantly D or G. Despite being variable, p15 shows biases toward P, S, T, L, E, K, Q and A. Using the relative aa frequencies and total size of the input database, we calculated the expected background levels of these octapeptides in randomized sequences (Fig. 4C, blue bars). Notwithstanding caveats about potential non-independence of the database sequences (see Methods), we flagged variants that occurred at more than three times the background level as probable functional variants. The most strongly over-represented were G, C or N at p12; well-supported examples of the rare C and N variations in a rotavirus polyprotein are shown in Fig. S3C-E. Thus, we expanded our core octapeptide motif to (D/G/C/N)(V/I)ExNPGP for further analyses (below). Other, rarer variants were T or V at p12, H at p14, H at p16 and M or W at p19.

**Figure 4.**
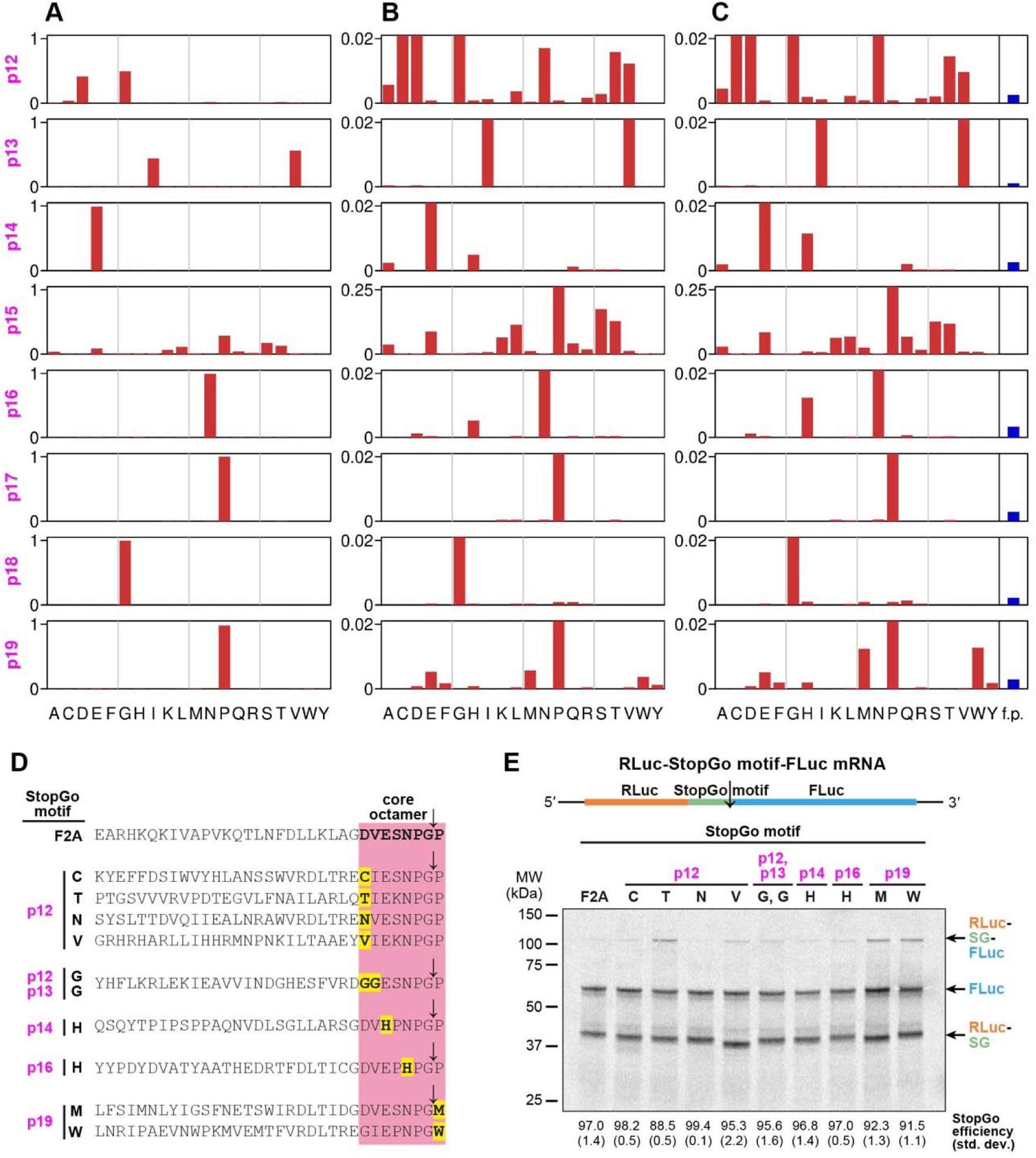
Bioinformatic identification of StopGo core octapeptide variants. Starting with the NCBI, Edgar, Neri, Zayed, Wolf and Wu sequences, ORFs were identified and clustered with CD-HIT using a 70% amino acid identity threshold. All octapeptides differing from D(V/I)ExNPGP by at most one position were identified, yielding 908 exact matches and 1557 inexact matches. **(A)** At each site, the proportion of the 2465 octapeptides exhibiting each of the 20 amino acids was quantified. **(B)** Same as panel A but with the y-axis enlarged 50x (4x for p15) to facilitate viewing of rare variants (the bars corresponding to prevalent variants extend beyond the top of the graphs). **(C)** Same as panel B but with amino acid proportions inversely scaled by overall amino acid frequencies in the input database. Blue bars at far right show the expected total number of octapeptides (i.e. summed over all possible amino acid substitutions) differing from D(V/I)ExNPGP at only the given position, in a randomised database with the same size and amino acid frequencies as the input database; f.p. denotes false positives. **(D, E)** StopGo sequences containing rare variants in the core octamer (D), used for the StopGo activity assay in (F) based on a dual reporter construct (top). The calculated StopGo efficiencies (%) and standard deviations (±%) are given below (*n* = 3).

Given that p12 = G is as prevalent as p12 = D, we performed the same analysis using octapeptides conforming to G(V/I)ExNPGP or differing at a single position (Fig. S4A-C). The results largely mirrored those from the D(V/I)ExNPGP set, confirming rare T/V (p12), H (p14, p16), and M/W (p19) substitutions. Additionally, we identified the rare variant p13 = G present exclusively when p12 = G.

Next, we evaluated the activity of the rare octapeptide variants using our dual reporter construct. We selected representative naturally occurring StopGo sequences containing the p12 = C/T/N/V, p13 = G, p14 = H, p16 = H or p19 = M/W variations (Fig. 4D), and compared their activity in RRL to the WT F2A construct (Fig. 4E). Intriguingly, all octapeptide variants showed activities of ∼90% or higher, i.e. close or equivalent to the F2A activity. This confirms the validity of our approach to search for functional StopGo variants using viral genomes and highlights the natural plasticity of StopGo sequences. Together, these findings provide a refined understanding of StopGo sequence flexibility.

The StopGo core octapeptide sequence is highly conserved at the amino acid level. To assess whether there are any constraints at the nucleic acid level, we analyzed codon usage at each position within the octapeptide across 176 and 98 StopGo sequences from chordate- and arthropod-infecting viruses, respectively, compared to codon usage in human and Drosophila coding sequences (Fig. S4D). Our analysis revealed no single dominant codon at any position within the StopGo octapeptide.

### Distinct sequence motifs exist upstream of the StopGo core octamer

Our structure shows clear density for at least 6 aa preceding the core octamer of the F2A-NC (Fig. 2A), that interact with the peptide exit tunnel and thus likely contributes to stabilizing the ribosome at the StopGo site (Fig 2B, C). Yet, it is well-known that “classical” StopGo motifs (e.g., F2A, E2A, P2A) differ in the positions upstream of the core octapeptide, despite the identity of specific amino acids in this region being important for StopGo activity (28). To investigate this upstream region, we returned to the 793,449 theoretical viral ORFs and extracted 100-aa regions flanking all occurrences of the expanded (D/G/C/N)(V/I)ExNPGP motif, yielding 9968 sequences. To select for evolutionary independence (see Methods), we clustered these at 50% identity (BLASTCLUST), and selected a representative sequence from each of the 1900 clusters (Fig. S3A).

A sequence logo (Fig. 5A) revealed non-random aa usage over six upstream residues (p6 to p11). If different upstream motifs are present in different subsets of sequences, then a sequence logo of all sequences combined may obscure such features. Therefore, to identify potential motif subgroups, we extracted the hexapeptide immediately upstream of the StopGo octapeptide and informally classified the 20 most frequent sequences into four major groups (PAPRLV-like, (R/K)DLTxE-like, DLTxDG-like, and LLLLSG-like) and one singleton (Fig. S3B).

**Figure 5.**
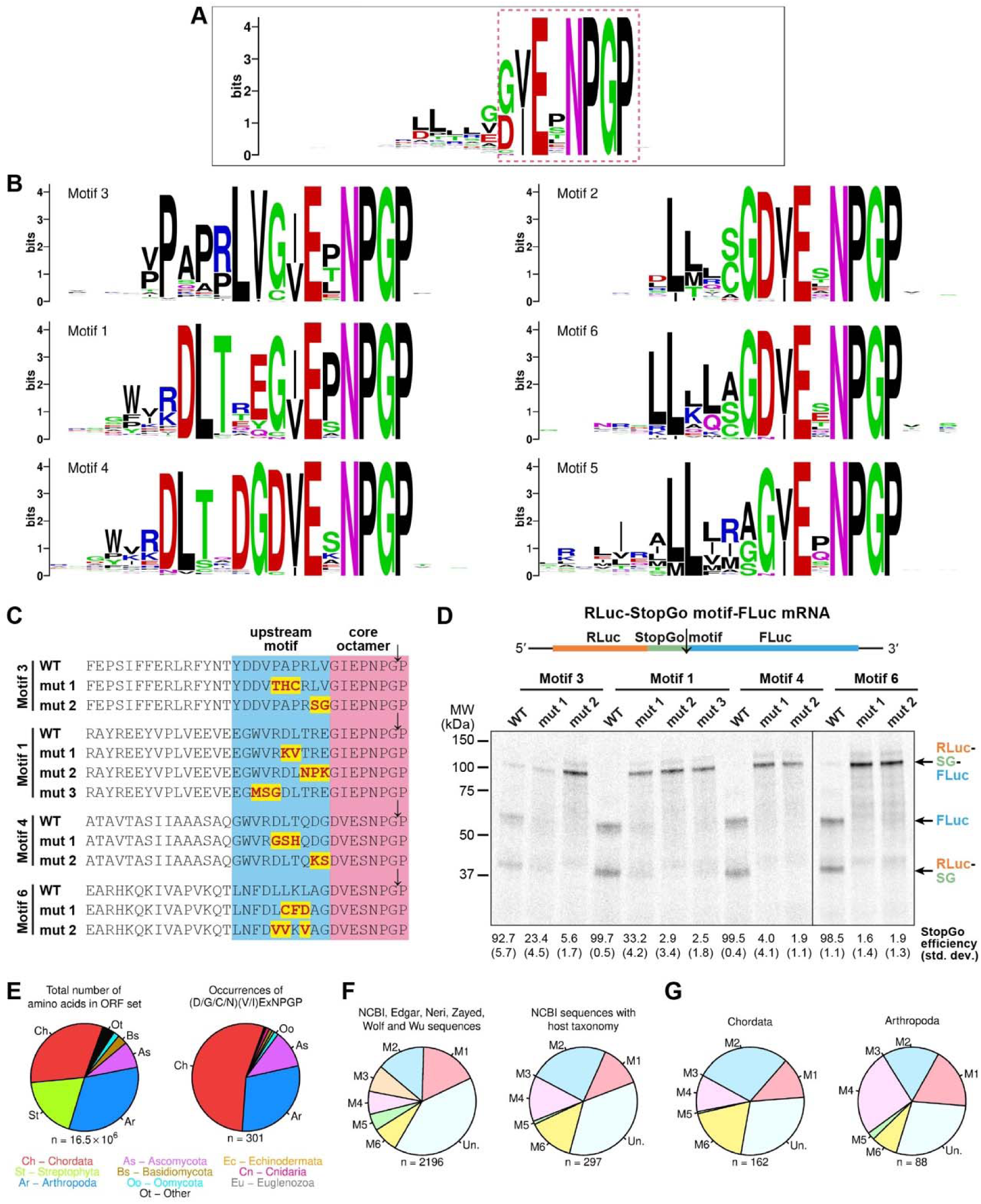
Identification of distinct motif classes upstream of the StopGo core octapeptide and host distribution of core octapeptides and upstream hexapeptide motifs. **(A)** Weblogo produced from 1900 representative StopGo sequences. The depicted region covers from 14 positions upstream of the octapeptide to 8 positions downstream. The core octapeptide is highlighted (red dashed box). **(B)** Weblogos for sequences identified by FIMO from among the 1900 representative sequences as matching each of the six MEME-defined hexapeptide motifs. **(C)** StopGo sequences containing examples of the upstream motifs identified in (B), and mutants of the most characteristic positions. **(D)** StopGo activity assay of the sequences in (D), based on a dual reporter construct (top). The calculated StopGo efficiency (%) and standard deviations (±%) are given below (*n* = 3). **(E)** Representation of host phyla in the input sequence database (total number of amino acids; left) and among the identified (D/G/C/N)(V/I)ExNPGP octapeptides (right) for NCBI RNA virus sequences with designated host species. **(F)** Representation of hexapeptide motifs 1–6 (M1 to M6), besides unclassified hexapeptides (Un.), upstream of (D/G/C/N)(V/I)ExNPGP octapeptides in the NCBI, Edgar, Neri, Zayed, Wolf and Wu sequences (left) or in the subset of NCBI RNA virus sequences with designated host species (right). **(G)** Representation of hexapeptide motifs upstream of (D/G/C/N)(V/I)ExNPGP octapeptides in NCBI RNA virus sequences with host species in phyla Chordata (left) or Arthropoda (right). In panels E–G, graphs show results for the reduced set of ORFs after CD-HIT clustering with a 70% amino acid identity threshold.

To explore upstream motifs in greater depth we used MEME (which identifies recurring motifs in sets of sequences) to look for the eight most statistically significant motifs among the 1900 hexapeptides (Fig. S5A). Motifs 7 and 8 were discarded due to low significance (E-values: 0.550 and 0.038). Notably, motifs 1, 3 and 4 correspond to the manually identified (R/K)DLTxE, PAPRLV and DLTxDG motifs, whereas motifs 2 and 6 share similarities with the LLLLSG motif. Using FIMO from the MEME suite, we mapped motifs 1–6 to hexapeptides immediately upstream of (D/G/C/N)(V/I)ExNPGP across our dataset. The 1900 representative sequences were grouped by motif (where found), and sequence logos were generated to visualize additional aa preferences in flanking regions (Fig. 5B). Some specific examples of usage of these motifs are shown in Fig. S5B-F.

Several trends are apparent. Motifs 1, 3 and 5 preferentially associate with p12 = G and p15 = P. In contrast, motifs 2, 4 and 6 associate with p12 = D and p15 = S. Interestingly, motifs 1 and 4 share a WV(R/K)DLTx[acidic]G signature, however they are offset by 1 aa, with the G corresponding to p12 in motif 1 but p11 in motif 4. Similarly, L-rich motifs 2 and 6 terminate in (A/S/C)G preceding p12, whereas motif 5 ends in (A/G/S)G at p12. A comparison of all 688 p12 = D sequences with all 1096 p12 = G sequences (Fig. S6A) revealed a strong G preference at p11 for the former but not for the latter.

To assess the contribution of upstream motifs to StopGo activity, we selected motifs 1, 3, 4 and 6 for in vitro activity assays in the RRL system, using the previously employed dual reporter construct with the F2A sequence replaced by a representative example of each motif (Fig. 5C,D). All four WT motifs supported StopGo activity; however, their efficiencies varied from ∼92.7% for motif 3 up to almost 100% for motifs 1 and 4. Mutating characteristic amino acids in each motif to amino acids that were rarely or never used in sequences matching the given motif led to a near- complete loss of StopGo activity in almost all cases. This highlights the essential role of upstream sequence context in enabling StopGo, suggesting that specific amino acids are required to engage functionally important interactions within the ribosomal peptide exit tunnel.

### Host distribution of StopGo sequences and the upstream motifs

To analyze the distribution of StopGo motifs across different host taxa, we used the database of 793,449 translated viral ORFs, besides a subset comprising NCBI sequences with designated host species (149,503 ORFs). After clustering at 70% aa identity (CD-HIT), we identified 131,941 and 12,688 representative sequences, respectively. We then searched all four datasets for occurrences of the expanded motif (D/G/C/N)(V/I)ExNPGP.

Among viruses with known hosts, StopGo sequences were predominantly found in chordate and arthropod viruses, in part reflecting the input dataset composition (Fig. 5E, S6C). However, despite plant viruses comprising 11% of input sequences, almost no StopGo motifs were detected in plant viruses. The six identified cases were in dicistroviruses and an iflavirus, both typically associated with arthropods, suggesting host misclassification. Interestingly, however, StopGo is known to function in artificial constructs expressed in plants (42) including in the context of plant virus vectors (e.g. (43)).

We also examined p12 residue distribution noting that, whereas G was more frequent in the full dataset, D was predominant in viruses of known host including chordate-, arthropod- and ascomycota-infecting viruses (Fig. S6B).

Analysis of the six upstream StopGo motifs revealed broad representation across datasets, though 28% of all sequences remained unclassified (Fig. 5F,S6D). Notably, however, motif 3 was absent from viruses with known hosts, being primarily detected in metagenomic datasets linked to environmental sources (e.g., soil, freshwater, sediment). In contrast, motifs 1, 2, 4 and 6 were common in chordate and arthropod viruses (Fig. 5G,S6E).

## Discussion

### Model of the StopGo mechanism

StopGo is a unique eukaryotic recoding phenomenon that allows co-translational separation of two proteins. Our structural studies of an F2A-bound ribosomal StopGo complex provide significant insights into the underlying mechanism and therefore allow us to update the model for StopGo (Fig. 6). In this model, when the ribosome encounters a StopGo sequence during elongation, interactions between the nascent polypeptide and the ribosomal exit tunnel induce initial ribosomal pausing. In particular, in the F2A case, the upstream L-rich motif forms a hydrophobic cluster that obstructs the exit tunnel, while the conserved D12, E14 and N16 residues establish hydrogen bonds with 28S rRNA, further prolonging ribosome pausing at the StopGo site. Consistent with this, primer extension assays demonstrated ribosome pausing with the G18 codon in the P-site and the P19 codon in the A-site (7) and ribosome profiling data on TMEV-infected cells indicated substantial disome formation at the StopGo site, indicative of ribosome collisions due to halted elongation complexes (44).

**Figure 6.**
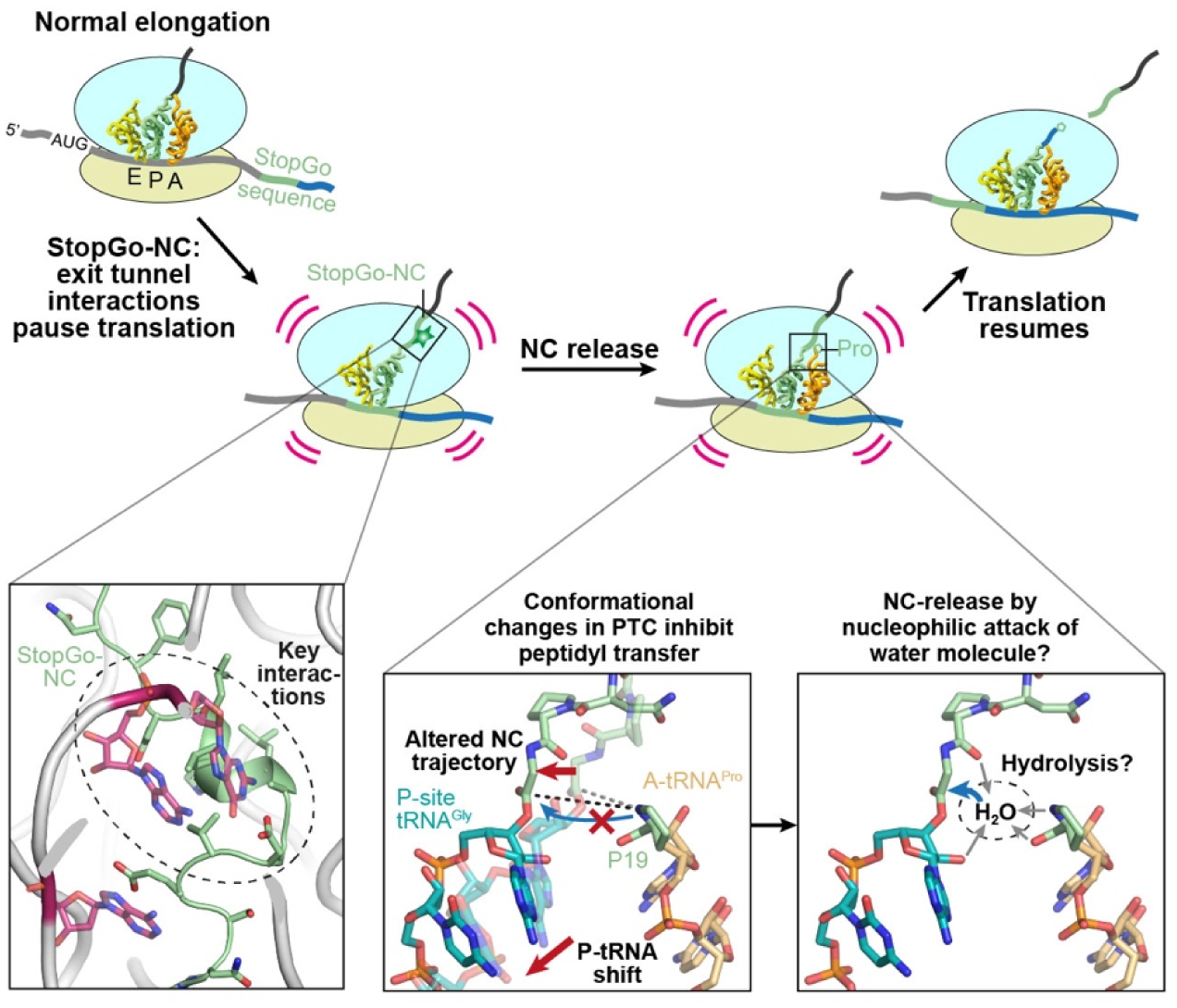
Proposed model for the StopGo mechanism. During translation of a StopGo sequence, the StopGo-NC (F2A in this study) interacts with the nascent peptide exit tunnel of the elongating ribosome, impeding ribosomal processivity (bottom left). A conformational change in the PTC induced by the StopGo-NC stabilizes the P-tRNA CCA end in an inactive state, which inhibits peptidyl transfer (bottom, middle) but may pre-expose the tRNA:StopGo-NC ester bond for hydrolysis (bottom right). After NC release, translation continues with the downstream product starting with P19. In the bottom middle panel, the F2A-NC was modeled in the canonical conformation based on PDB ID 7O80 (transparent P-tRNA/NC); the A-tRNA^Pro^ was modeled based on PDB ID 6Y0G.

Interactions between the F2A-NC and the ribosomal exit tunnel facilitate the formation of a distinct loop structure by the highly conserved NPG motif. Notably, P17, with its cyclic side chain restricting φ and ψ dihedral angles, adopts a conformation opposite to N16. Together with G18, which due to lack of a side chain can sample a large range of the φ and ψ angles, this enables the residues to adopt the specific conformation that induces a structural rearrangement which pushes the P-tRNA^Gly^ CCA end outwards, away from the exit tunnel (Fig. 3A-C). Concurrently, this triggers conformational changes in the PTC, most notably the swapping of the U4198 sidechain from a position that laterally lines the NC towards the A-site, to a position above the NC. This makes its conformation distinct from all other previously solved structures of both the uninduced and induced states. Overall, the conformational changes move the P-tRNA CCA end away from its “canonical” position required for addition of the next amino acid. This renders the elongation complex into an inactive state, minimizing the likelihood of peptide bond formation by the incoming A-tRNA and prolonging the pause at the StopGo site. It is likely that the CCA end and C-terminus of the F2A-NC just after addition of G18 to P17 is in the “canonical” conformation of a catalytically active elongation complex. Until the characteristic F2A-NC conformation and the CCA/PTC rearrangements are fully formed, this leaves a temporal window for addition of the next amino acid (P19), which would bypass the StopGo event (i.e. allowing uninterrupted translation). We speculate that one function of the upstream residues is to slow down the elongating ribosome and provide a window for the rearrangements to occur.

Recent structures of bacterial termination complexes have suggested an updated model for NC release during canonical termination. This model proposes an RF induced switch of the A76 sugar pucker from the usual C2′ endo to a C3′ endo conformation. The 2′ OH group may then be positioned to act as a nucleophile for an attack on the ester bond (Fig. S7A). The resulting 2′, 3′ cyclic intermediate could subsequently be hydrolyzed by “bulk” water for NC release (45). In our structure, the A76 ribose is in C2′-endo conformation (Fig. S7A), similar to the pre- transpeptidation state, and the 2′ OH group is positioned towards, but not directly above the ester bond (Fig. S7B). Compared to the termination complex, both the distance of the 2′ OH group to the ester CO carbon (3.6 Å vs 3.7 Å) and the BD angle (2′ O-C^O^-O, 112° in both cases) are nearly identical, however, the angle of the 2′ OH group towards the carbonyl plane is 10° more shallow in our structure (Fig. S7C). Therefore, we conclude that a 2′ OH-initiated release reaction, analogous to canonical translation termination is unlikely. How could NC release in the StopGo case happen? Unlike our eRF1-AAQ stalled complex, a native StopGo complex would contain tRNA^Pro^ bound to the A-loop, which would stabilize the PTC in the induced state. This would result in movement of U4277 (U2585), which would then even further expose the ester bond, that is already partially uncovered from its shielded position due to the conformation imposed by the NC. Modelling an A-site tRNA^Pro^ in place of eRF1-AAQ shows that this would result in a gap that might allow coordination of a water molecule above the ester plane through e.g. the P-site A76 2′ OH and the Pro carboxyl/amino groups (Fig. S7D). The role of the incoming tRNA would therefore consist of fully uncovering the ester bond and providing a scaffold for binding an “activated” water. However, follow-up work will be required to determine the precise mechanism of NC hydrolysis.

After the StopGo-NC has been hydrolysed and dissociated through the exit tunnel, the P-tRNA and PTC most likely reset to their “canonical” positions. The ribosome then contains an acylated A-tRNA and a deacylated P-tRNA – a situation homologous to the product state of a regular peptide bond formation event – and will then transition through the hybrid state that eventually allows for eEF2 mediated translocation and normal continuation of translation elongation.

### The diversity of StopGo/2A recoding

StopGo is commonly used by RNA viruses but only rarely in cellular genes (12, 13, 17, 18). The reason for this may be, in part, due to the much higher substitution rates of RNA viruses allowing them to rapidly explore sequence space and evolve new functions. By searching RNA virus datasets for StopGo-like octapeptides occurring more frequently than random, we inferred the existence of functional rare variants. These all proved to be highly active when tested within naturally occuring flanking sequence contexts. Variants p12 = C/N/V, p13 = G, p14 = H and p16 = H displayed >95% activity; p19 = M/W had 92.3/91.5% activity; and p12 = T had 88.5% activity. Interestingly, the p19 = M/W variants exhibited considerably higher activities than corresponding p19 mutants in the F2A context (92.3/91.5% versus 55.3/80.5%). This may indicate an interplay between the formation of the release-competent, specific NC conformation (primarily provided by the residues up until p18) and the release step (p19). Similarly, D12N and N16H mutants had previously been found to be only modestly active when tested in the foreign context of EMCV 2A or F2A, respectively (30, 46, 47). Several other variants have been found to be partially active in earlier studies using mutated sequences (4, 10, 28, 30, 48) and in the present work (Fig. 3E) but were not flagged by our bioinformatic analysis. Part of the reason may be the high threshold that we used in the analysis – for example an E14Q mutation was previously found to permit 56% or 26% activity (10, 48) and in our analysis this variation was apparent in Fig. 4C albeit below threshold.

That the rare variants generally mediate efficient separation raises the question of why these variants are rare. One possibility is that the flanking sequence contexts that allow rare variants to function efficiently are more restrictive than for the canonical octapeptide. In cases where separation is less efficient (p12 = T, p19 = M/W), this may be a due to differences between RRL ribosomes and the ribosomes of the natural host organisms (with nearly all examples coming from viruses of unknown host). It is also possible that <100% separation could be functionally important in certain situations. Indeed one likely example of this arises from the presence of a StopGo site following an N-terminal signal sequence in multiple proteins in the sea urchin *Strongylocentrotus purpuratus*: when StopGo separation occurs the downstream product is cytoplasmic and when StopGo separation fails the full-length product is targeted to the exocytic pathway (49). Although it is tempting to speculate that rare variants may be selected in cases where less efficient separation is beneficial, StopGo sequences with canonical octapeptides can also work in this way (12, 49).

A codon usage analysis revealed no evidence for any stimulator in the nucleotide sequence, consistent with previous findings (9, 11, 15) and consistent with the canonical positioning observed for the mRNA in our cryo-EM structure. Although differences between StopGo and host codon usage were apparent in our analysis (Fig. S4D), and StopGo codon biases have also been commented on elsewhere (47), these could arise from other factors such as dicodon biases, statistical noise, global differences between virus and host codon usage, and selection against CpG dinucleotides in many vertebrate viruses due to the antiviral action of the interferon-stimulated ZAP protein (50).

By applying MEME and FIMO analyses, we identified several distinct upstream motifs associated with the core StopGo octapeptide. Although we labelled six motifs, the actual number is not fixed: motifs can be merged into fewer groups (e.g. the L-rich motifs 2 and 6), or additional motif groups can be defined (e.g. the cardiovirus-like p11 = H motif) depending on the parameters and depth of the analysis. Furthermore, 28% of our sequences remained unclassified. As noted previously (28), it appears that there are multiple different ways in which the upstream sequence can assist in positioning the core octapeptide to promote StopGo activity. Notably, chimeric sequences are often inactive (28).

We defined our motifs based on the six amino acid positions preceding the core octapeptide. However, it is worth re-emphasising that StopGo activity depends on more than just 14 aa, with full activity in FMDV requiring >30 aa of native sequence (5, 29). Although early work suggested a propensity for the upstream sequence to form an alpha helical structure (4), later work reported this not to be a universal feature of StopGo (12, 28). An analysis of alpha helical propensity by motif type (Fig. S8) confirmed this, with motif 3 lacking predicted alpha helical propensity. Motifs 1, 3 and 5 associate with p12 = G, whereas motifs 2, 4 and 6 associate with p12 = D. In the F2A- StopGo complex, D12 forms two hydrogen bonds with rRNA, and its substitution with Ala abolishes StopGo activity, highlighting the critical role of its side chain. However, motifs 1, 3 and 5 maintain StopGo function despite having G at p12, likely requiring compensatory interactions from upstream residues.

Usage of motifs 1, 2, 4 and 6 was widespread among chordate and arthropod viruses but, surprisingly, motif 3 was only found in virus sequences derived from metagenomic datasets and never in a virus of known host. Although StopGo activity of a motif 3 sequence was verified (Fig. 5D) it was somewhat inefficient in RRL, consistent with it never being observed in mammalian viruses. Motif 3 differed substantially from the other motifs, being proline rich and lacking alpha helical propensity. Another interesting feature is that there is a clear association between upstream motifs and the aa preferences at the p12 and p15 positions in the octapeptide. Further, motifs 1 and 4 are strikingly similar but offset by 1 aa, indicating that NC “tension” is unlikely to be relevant for StopGo activity, but consistent with NC amino acids making specific interactions with the exit tunnel.

In conclusion, our study provides structural and sequence-level insights into the StopGo mechanism, revealing how the interactions between the StopGo-NC and the ribosomal exit tunnel induce conformational changes in the PTC, that lever the tRNA:StopGo-NC ester bond out of its protected position to promote peptide release without ribosome disassembly. A large-scale database search confirmed that StopGo octapeptides are widespread within RNA viruses. As mutating StopGo sequences strongly impacts viral fitness (47), but StopGo is never or only rarely used in cellular organisms (13), this presents an opportunity to develop inhibitors for therapeutic intervention in the future, e.g. by structure-guided design. Further analysis revealed rare variants that we found to be active in vitro. Especially interesting for application purposes is the p19 = M variant, which would allow the downstream encoded protein to begin with Met, therefore avoiding potential destabilizing effects of N-terminal Pro residues (51). Overall, our results therefore provide a unified model of the StopGo mechanism and set the stage for improved engineering of StopGo sequences as valuable tools in biotechnology and utilization of StopGo as a potential target for antiviral drugs.

## Acknowledgments

We thank Dr Y. Gordiyenko for preparation of eRF1-AAQ, and Dr V. Chandrasekaran and Dr Y. Du for helpful discussions and advice in cryo-EM data processing. We also thank the Electron Microscopy and Scientific Computing facilities of the MRC Laboratory of Molecular Biology for access to and support of electron microscopy and computing clusters, respectively. X.L. was supported by an EMBO Postdoctoral Fellowship (ALTF 937-2022). P.K.Z. held Postdoctoral fellowships from the German National Academy of Sciences Leopoldina (LPDS 2021-14) and EMBO (ALTF 778-2021). This work was supported by a UK Medical Research Council grant (MC_U105184332) and the Wellcome Trust Senior Investigator award (WT096570) to V.R., the Irish Research Council Advanced Laureate Award (IRCLA/2019/74) to J.F.A., and a Wellcome Trust Senior Research Fellowship 220814/Z/20/Z to A.E.F. For the purpose of open access, the authors have applied a CC BY public copyright license to any Author Accepted article version arising from this submission.

## Materials and Methods

Plasmids encoding templates for cryoEM and StopGo activity assays were cloned using standard methods and mRNAs were subsequently transcribed using T7 RNA polymerase. StopGo activity assays were performed in RRL containing ^35^S-Met for detection via SDS-PAGE and autoradiography. Complexes for cryoEM were prepared by in vitro translation of the F2A-PGX mRNA in RRL supplemented with eRF1-AAQ, and subsequently purified via FLAG affinity chromatography. CryoEM data were collected on a Titan Krios microscope and then processed in RELION; a molecular model was build using Coot and refined in Phenix. Viral StopGo core octapeptide sequences were searched in the NCBI nr/nt database and other published datasets (37-41). Upstream motifs were identified using fimo (52) from the MEME package. For more information on construct design and molecular cloning, preparation of mRNAs, StopGo activity assays, cryoEM sample preparation, data collection, processing and model building, and bioinformatic analysis please consult the SI Appendix.

