## Supplemental Information for "Molecular architecture and diversity of StopGo/2A translational recoding"

**This PDF file includes:**

Supporting Information text
Materials and Methods
Figures S1 to S8
Table S1
SI References

### Supporting Information text

#### Comparison of the strategy for identification of StopGo motifs presented here to Rao et al. (2025)

A recently published report by Rao and colleagues (1) used a profile Hidden Markov Model (pHMM)-based approach to identify potential StopGo sequences, in contrast to our octapeptide-based search. We focused on RNA virus proteomes whereas Rao et al searched both virus and cellular proteomes. We separately analysed rare variants in the core octapeptide and defined motif groups based, initially, on the upstream hexapeptide, yielding six main motif groups. In contrast, Rao et al incorporated both into a single analysis (the pHMM models) and reported two main motif classes, A and B, that are broadly similar to our motifs 6 and 2, and motifs 1 and 4, respectively. In terms of taxonomic distribution of StopGo usage, we assessed the hosts of viruses containing different StopGo motifs whereas Rao et al analysed the distribution of StopGo motifs in virus and cellular taxa. An advantage of the former approach is that, because StopGo is used only rarely in cellular genes, its absence cannot be used as an indicator of the functionality of StopGo motifs in a given taxonomic group (e.g. StopGo is apparently absent from mammalian genomes, but frequently utilized in mammalian viruses). An advantage of the latter approach is that many cellular taxa (e.g. members of the SAR supergroup) are poorly represented in our database of viruses with known hosts.

### Materials and Methods

#### Construct design and molecular cloning

For cryo-EM studies of F2A-mediated StopGo, we generated the F2A PGX in vitro transcription construct (Fig. 1A) based on the FMDV serotype SAT-2 genome/polyprotein sequences (accession numbers: NCBI accession AJ251473, UniProt accession Q9Q2N9). The construct encodes the C-terminus of VP1 (K183-Q214), all of 2A, and the N-terminus of 2B (P1-L14), corresponding to residues K908-L971 of the FMDV polyprotein. Position 1 of 2B (i.e. p19 of F2A) was changed to a TGA stop codon to enable stable stalling of the target complex (making the 2B region the 3′ UTR of the corresponding mRNA). The FMDV sequence was N-terminally fused to a 3X-FLAG tag, a PreScission protease cleavage site and a 29 aa long synthetic linker. To enable efficient translation in RRL, a short (64 nt) 5′ UTR containing CAA repeats at the 5′ end, preceded by a T7 promoter was added upstream to the coding sequence. The construct was synthesized as a gene block by IDT and used as template for PCR (see below).

Plasmid pDLuc (2) was used for StopGo activity assays. The plasmid encodes a single ORF composed of an N-terminal ("upstream") Renilla Luciferase (RLuc) and a C-terminal ("downstream") Firefly Luciferase (FLuc). The ORF is preceded by a 37 nt 5′ UTR and followed by a 153 nt 3′ UTR; transcription is driven from a T7 promoter. Although a luciferase reporter construct, herein it was used simply as a reporter in radiolabelled SDS-PAGE analysis.

To test the activities of rare variants of the StopGo core octapeptide (Fig. 4D, E), representative StopGo sequences (33 aa up to and including p19), with p12 = C (human rotavirus B219; NCBI accession DQ168032), p12 = N (porcine rotavirus H; NCBI accession LC348489), p12 = T (ND_130348), p12 = V (SRR6823499.macro.NODE_1292_length_3293_cov_3.084472), p13 = G (ERR1356705.macro.NODE_468_length_3723_cov_32.322265), p14 = H (ND_032879), p16 = H (ND_051457), p19 = M (SRR8554514.macro.NODE_40_length_9054_cov_28.836210) or p19 = W (ERR2562196.macro.NODE_10_length_9841_cov_104.213452) ("ND" accessions correspond to the RiboV1.4_Contigs.fasta file of (3), "SRR"/"ERR" accessions correspond to the rdrp_contigs.fa file of (4)) were reverse translated, and then codon-optimized for mammalian (human) expression to exclude unwanted codon-level effects during in vitro translation in RRL. The resulting sequences were then synthesized as genes blocks (Azenta) and subsequently inserted between the FLuc and RLuc ORFs in pDLuc via Gibson assembly.

For evaluation of the influence of upstream motifs on StopGo efficiency (Fig. 5D), codon-optimized versions of representative sequences (33 aa up to and including p19) for motifs 1 and 4 (both from Lanama virus; NCBI accession MW218667), motif 3 (from SRR12063586.macro.NODE_24_length_9085_cov_41.544326; rdrp_contigs.fa file of (4)) and motif 6 (FMDV serotype O; NCBI accession X00871) were inserted between FLuc and RLuc in pDLuc via PCR using diverging 5′ phosphorylated primers and subsequent DNA ligation. Upstream motif mutants were designed to exchange characteristic amino acids of each motif for amino acids that were rarely or never used at the corresponding positions in sequences matching the given motif (Fig. 5C).

For analysis of the importance of p19 on StopGo efficiency, the pDLuc vector containing the FMDV StopGo sequence was used as a basis for mutation to all other 19 amino acids using the QuickChange protocol (Agilent Technologies).

All molecular cloning was carried out using standard methods, including Q5 DNA polymerase (NEB, cat. #M0491) driven PCR, Gibson assembly using NEBuilder HiFi DNA assembly master mix (NEB, cat. #E2621), T4 DNA ligase (NEB, cat. #M0202) for ligation and *E. coli* TOP10 cells for heat-shock based transformation. Constructs were verified by Sanger sequencing.

### Preparation of mRNAs

For generation of mRNA for cryoEM, the gene block encoding the F2A PGX construct was PCR amplified to generate a template for in vitro transcription using the Ampliscribe T7 Flash system (Lucigen). The mRNA was then purified by lithium chloride precipitation and re-dissolved in $H_2O$. All mRNAs for StopGo activity assays were generated using T7 RNA polymerase driven in vitro transcription. Templates were prepared via PCR (Q5 DNA polymerase; NEB, cat. #M0491), *Dpn*I-digested (NEB, #R0176), and subsequently purified using a QIAquick PCR purification kit (Qiagen). The PCR products were then used at 20–25 ng/µl in a 100 µl reaction mix containing 80 mM HEPES/KOH (pH 7.5), 24 mM $MgCl_2$, 2 mM spermidine, 40 mM dithiothreitol (DTT), 6 mM each of ATP, GTP, CTP and UTP, 1 U/ml RNaseIN and 14 µg/ml of home-made T7 RNA polymerase. In vitro transcription was carried out for 4 h at 37 °C, then the templates were digested by addition of 1 µl of RNase-free DNase I (NEB, cat. #M0303) and further incubated for 30 min at 37 °C. The RNAs were then lithium chloride precipitated overnight at −20 °C, the pellets collected by centrifugation, washed with 70% ethanol, and the mRNAs finally dissolved in $H_2O$ at concentration of ~1.4 µg/µl.

### StopGo activity assays

Activity of the StopGo sequences in the pDLuc plasmids was assessed by in vitro translation using a rabbit reticulocyte lysate (RRL; Green Hectares, USA) based system (5). Reactions typically contained 0.5 volumes of micrococcal nuclease-treated, supplemented RRL (“cT2”), 10 µM of Amino Acid Mixture Minus Methionine (Promega, cat. #L996A), 0.5 U/µl RNaseIN (Promega, cat. #N251A) and 0.5 µCi/µl EasyTag L-35S-Methionine (Revvity, cat. #NEG709A500UC) and 25 ng/µl of mRNA in 8 µl total reaction volume. The in vitro translation was carried out for 25 min at 32 °C, then 2 µl of 0.25 mg/ml RNase A was added. After 5 min incubation at room temperature, 5 µl of 4x LDS loading dye was added and 3 µl of sample was subjected to SDS-PAGE (NuPAGE Bis-Tris gradient gels, 4–12%). Gels were dried using a slab gel dryer and then used to expose a BAS-IP SR 2025 Imaging plate (Amersham). Autoradiograms were visualized after 1–2 days of exposure using a Typhoon Imager (Cytiva). All in vitro translation experiments were performed at least as triplicates.

Data were analysed using the Gel Analysis tool in ImageJ v. 1.53k (6). A background correction using a “rolling ball” with radius 65 was applied before integration of the signals. To calculate the StopGo efficiency, the signals of RLuc, FLuc and the RLuc-FLuc fusion product resulting from unsuccessful StopGo events, were normalized to the number of Met residues in the corresponding polypeptides. Efficiencies were calculated individually for RLuc and FLuc as the ratio of normalized signals of RLuc or FLuc, respectively, divided by the sum of normalized RLuc or FLuc and the RLuc-FLuc fusion product signals. The resulting individual efficiencies were then averaged.

### Cryo-EM sample preparation and data acquisition

Translationally active RRL was generated from untreated rabbit reticulocyte lysate (Green Hectares, USA) as described in (5) with certain modifications. The final translation reaction (per 20 µl) contained 11.4 µl nuclease-treated and activated RRL, 200 ng/µl mRNA, 1.2 mM $MgCl_2$, 0.5 mM DTT, 0.4 mM spermidine and the AAQ variant of human eRF1 (eRF1-AAQ; purified as described earlier (7)) at a final concentration of 3.5 µM to trap translating ribosomes. Reactions were incubated for 25 min at 32 °C. 4 ml translation reactions were chilled on ice for 10 min to halt translation, and HEPES/KOH (pH 7.5) was added to a final concentration of 50 mM. Chilled translation reactions were directly incubated with 400 µl of packed anti-FLAG M2 beads (Sigma, cat. #A2220) for 2 h at 4 °C with gentle mixing. The beads were then washed with 4 ml of 50 mM HEPES (pH 7.4), 100 mM KOAc, 5 mM $Mg(OAc)_2$, 0.1% Triton X-100, 1 mM DTT; 4 ml of 50 mM HEPES (pH 7.4), 250 mM KOAc, 5 mM $Mg(OAc)_2$, 0.5% Triton X-100, 1 mM DTT and 6 ml of RNC buffer (50 mM HEPES (pH 7.4), 100 mM KOAc, 5 mM $Mg(OAc)_2$, 1 mM DTT). Ribosomes were eluted after 4 sequential 10 min incubations at room temperature in RNC buffer that contained 0.2 mg/ml 3x FLAG peptide (Sigma, cat. #F4799). The elutions were combined and centrifuged at 186,000 x g at 4 °C for 2 h in a TLA 55 rotor (Beckman Coulter). Supernatant was discarded and the pellet was resuspended in RNC buffer at a concentration of 100 nM. At each step of translation reactions and purification, aliquots were taken to perform SDS-PAGE on 12 % polyacrylamide gels and Western blots. Gene products of interest were probed using anti-FLAG antibody (Sigma Cat. No. A8592).

For grid preparation, QUANTIFOIL Au 300 mesh, R1.2/1.3 grids (Quantifoil) were glow-discharged for 45 s using an Edwards glow discharger operated at 0.1 torr and 30 mA. The grids were then modified by application of 3.5 µl of 0.2 mg/ml graphene oxide solution (Sigma, cat. #763705) for 1 min. The solution was then blotted off, the grids washed three times with 20 µl $H_2O$ and finally dried for 1.5 h before use. Grids were frozen using VitroBot Mark II (Thermo Fisher Scientific) operated at 4 °C and 100% relative humidity. 3.5 µl of sample were applied to the graphene oxide coated side of the grids and incubated for 30 s to allow adsorption and concentration of ribosomes. The sample was then blotted off for 3.5–4.5 s at a blot force of −3 using a double layered Whatman 595 filter paper before rapid plunging into liquid ethane cooled to −170 °C using a temperature-controlled thermostat. Grids were then clipped and stored in liquid nitrogen until data collection.
Cryo-EM data were collected on a Titan Krios G1 microscope (Thermo Fisher Scientific) at the LMB EM facility. The microscope was equipped with a Gatan Quantum energy-filter and a Gatan K3 post-GIF direct electron detector operated in counting mode. Data were collected using EPU software in faster acquisition setting (AFIS), using a defocus range of −1.0 to −2.2 µm at 105,000x magnification, corresponding to a nominal pixel size of 0.725 Å. The total dose per micrograph was 40 $e^-/Å^2$, distributed over 40 fractions.

### Data processing

Cryo-EM data were processed using RELION v. 4.0 and 5.0 (8). Raw movies were first motion-corrected using MotionCor2 (9), then the contrast transfer function (CTF) was estimated with CTFFIND v. 4.1 (10). Micrographs with a resolution worse than 5 Å were removed. Particles were picked using template matching of 80S ribosome 2D classes obtained from manual picking and 2D classification. The particles were then extracted, subjected to initial 3D classification for identification of true ribosomal particles (reference map: EMD-0194) and further processed by (i) 3D classification without image alignment, (ii) focussed classification for eRF1/eRF1-AAQ, and (iii) focussed classification on the P-site tRNA:NC with signal subtraction. The complete processing pipeline is depicted in Fig. S1B. The pixel size of the final map (0.702 Å/pixel) was calibrated in Postprocessing step using a model of the eRF1-AAQ bound 80S ribosome (PDB-ID 7O80). Local resolution (Fig. S1E) was estimated using RELION's implementation.

### Model building refinement and validation

The model of the eRF1-AAQ-stalled rabbit ribosome at the SARS-CoV-2 slippery site (PDB-ID 7O80, (11)) was docked into the final postprocessed F2A-StopGo map in Chimera v. 1.16 (Pettersen *et al*, 2004). The model was then further modified in Coot v. 0.9.8.94 (12) to model the F2A-NC, mRNA sequence and P- and E-site tRNAs. The model of the P-site tRNA-$Gly^{CCC}$ was based on a crystal structure of human tRNA-Gly (PDB-ID 4KR3), and the model for the E-site tRNA-$Pro^{AGG}$ was based on the corresponding E-site tRNA in PDB-ID 5LZV. Both tRNAs were mutated to reflect the sequence of rabbit tRNAs, as obtained from the GtRNAdb (13). The density was then inspected and compared to a list of common tRNA modifications (14); where clearly identifiable in the density, the appropriate modification was then added. The tRNAs were numbered according to the established conventions (14). The mRNA density matched the corresponding sequence StopGo site and was adjusted accordingly. The F2A-NC was modelled starting at the C-terminus up to the point where the register could not be clearly matched any more to the corresponding density at the N-terminal end. This allowed placement of the entire FMDV 2A peptide and of six preceding amnio acids (i.e. I209–Q214 of VP1). Numbering of the NC reflects the numbering of the FMDV 2A protein. The model was subsequently refined using phenix.real_space_refine in Phenix v. 1.21 (15). A custom distance restraint (1.4 Å) and 120 degree bond angle restraints between the tRNA-Gly A76 3′ OH and the carboxyl group of the G18 residue of the NC were included to reflect the corresponding ester bond. Additionally, a nonbonded weight of 1000 was used. The resulting model was then further iteratively inspected in Coot to fit any outlying residues, correct geometry and place ligands ($Mg^{2+}$, $K^+$, spermidine) and re-subjected to refinement in Phenix. The final model was validated, and half map/model-map Fourier shell correlation (FSC) curves were calculated using the comprehensive validation tool in Phenix.

**Bioinformatic analysis of StopGo sequence diversity**

RNA virus and retrovirus (realm Riboviria) nucleotide sequences in the length range 1500 to 50,000 nt were identified in the NCBI nr/nt database using the search query txid2559587[Organism:exp] AND 1500:50000[Sequence Length], and downloaded between 17 and 20 Feb 2023. To reduce download time, seven taxa with extremely large numbers of sequences (severe acute respiratory syndrome-related coronavirus, human immunodeficiency virus 1, influenza A virus, hepacivirus C, hepatitis B virus, influenza B virus and rotavirus A; NCBI taxonomy IDs 694009, 11676, 11320, 11103, 10407, 11520 and 28875, respectively) were excluded, and instead only the NCBI RefSeq sequences (where ≥1500 nt) were downloaded for each of these. Patent and synthetic sequence records were removed; this is important since, for example, the identification of 2A peptides in virus phylum Negarnaviricota reported in (1) actually results from synthetic sequences KR781609, MK271062, MT227009 and MK672825, where a foreign 2A cassette was artificially inserted into a virus sequence. Sequences with >10 ambiguous nucleotide codes (for example "N"s), indicative of low quality or incomplete sequencing, were also removed. This NCBI dataset was then combined with RNA virus nucleotide sequences from (3, 4, 16-18), again selecting only sequences in the length range 1500 to 50,000 nt, and removing any sequences with >10 ambiguous nucleotide codes. The resulting dataset comprised 1,056,829 sequences: 253,968 from NCBI and 527,294, 260,928, 4242, 1191 and 9206 from the Edgar, Neri, Wolf, Wu and Zayed datasets, respectively.

To define a subset of sequences with known host species taxonomy, we used NCBI sequences where the first two words in the “/host” tag in the GenBank record comprised a valid species Latin binomial (and not a generic term such as “dog” or “Canis sp.”). We used the names.dmp and nodes.dmp files from https://ftp.ncbi.nlm.nih.gov/pub/taxonomy/taxdump.tar.gz (downloaded 23 Oct 2024) to determine phylum names for each host species name. The resulting dataset comprised 147,544 NCBI sequences with assigned host phylum.

To remove identical sequences (or subsequences) the sets of 1,056,829 and 147,544 nucleotide sequences were each clustered with cd-hit-est (19) at the 100% identity threshold (options −c 1.00 −n 11 −M 50000 −d 0 −T 7 −g 1), with the longest sequence being retained as the representative in each cluster. This resulted in 869,685 and 126,979 representative nucleotide sequences, respectively. Start-codon-to-stop-codon ORFs, on either strand, with length ≥1500 nt were identifed with EMBOSS getorf (20) using the standard genetic code, yielding 793,449 and 149,503 ORFs, respectively. ORF amino acid sequences were clustered with cd-hit (19) at the 70% identity threshold (options −c 0.70 −n 5 −M 50000 −d 0 −T 7 −g 1), with the longest sequence being retained as the representative in each cluster. This resulted in 131,941 and 12,688 representative amino acid sequences, respectively. The four sets of translated ORFs contained respectively 10,266 (all ORFs), 1172 (ORFs with defined host taxonomy), 2242 (cd-hit 70%) and 301 (cd-hit 70%, defined host taxonomy) exact matches to (D/G/C/N)(V/I)ExNPGP.

To investigate sequences flanking the StopGo core octapeptide, we only used the exact matches to (D/G/C/N)(V/I)ExNPGP where the corresponding ORF encoded at least 42 aa upstream and at least 50 aa downstream of the octapeptide, resulting in 9968, 1151, 2196 and 297 instances, respectively. Using the 9968 instances (i.e. from all ORFs, without cd-hit 70% clustering), the 100 aa flanking sequences were extracted and clustered with BLASTCLUST (21) with options −a 8 −p

T −L 0.8 −b T and identity thresholds ranging from 40% to 100% in steps of 5% (Fig. S3A). We chose a relatively low identity threshold (viz. 50%; 1900 clusters) so that the resulting examples of StopGo would be more likely to be evolutionarily independent or, if evolutionarily related, amino acid sequences unrelated to StopGo activity (e.g. due to homology of the upstream or downstream protein domains) would be more likely to have accumulated differences. A representative 100 aa sequence was chosen from each cluster – either the most duplicated sequence (with ties broken by taking the alphanumerically first accession) or, if no 100 aa sequence was duplicated, the centroid sequence (minimum summed pairwise amino acid identity distances from sequence *i* to all other sequences *j* within the cluster).

To identify overrepresented motifs within the hexapeptide immediately upstream of the core octapeptide within the 1900 representative sequences, we used meme (22) from the MEME package ((23); version meme-5.3.3). We applied meme to the 1900 hexapeptides, with options −protein −mod zoops −nmotifs 8 −w 6 −objfun de −markov_order 0 −p 8 −neg negative.fa (i.e. each input sequence may contain 0 or 1 occurrences of a motif, and MEME will look for the eight most statistically significant motifs of width six amino acids). The file negative.fa is a negative control set of sequences that are expected to not contain StopGo-upstream-adjacent hexapeptides. To generate this file, we extracted the 50 aa immediately downstream of the core octapeptide for the 1900 representative sequences, removed 37 sequences that contained an additional NPGP in this region (to avoid having a StopGo in the negative control in cases of closely spaced StopGo sequences), and then removed the C-terminal 10 aa in case of additional NPGPs just downstream of the extracted 50 aa, resulting in 1863 sequences of length 40 aa. A total of 900 of the 1900 sequences were selected by meme to form motifs 1–6 (Fig. S5A).

Next, we used fimo (24) from the MEME package, with option −−thresh 1.0E−4, to search for motifs 1–6 in the hexapeptide immediately upstream of the core StopGo octapeptide for the 1900 representative sequences, and also for the StopGo octapeptides found in each of the four sets of ORFs (i.e. with or without defined host taxonomy, with or without cd-hit 70% clustering). Where a hexapeptide was matched to more than one of motifs 1–6, we assigned it to the motif with the best fimo *p*-value. Among the 1900 sequences, we assigned 334, 243, 175, 120, 105 and 102 to motifs 1–6, respectively, with the remaining 821 being unclassified. While the fimo assignments did not agree precisely with the original meme assignments, this system allowed us to extend identification of motifs 1–6 beyond the 1900 evolutionarily diverse representatives that were used to define the motifs. The fimo-annotated motif-match sets tend to be larger than those used by meme to initially generate the motifs, so the resulting logos in Fig. 5B are slightly different from the meme logos in Fig. S5A.

Sequence logo figures were produced via the Weblogo server at https://weblogo.berkeley.edu/logo.cgi.
The rdrp_contigs.fa file (version 27, Jun 2021) containing the (4) sequences was downloaded from https://github.com/ababaian/serratus/wiki/Viral-contigs-containing-RdRP.

The expected number of occurrences (in a randomized database of the same size as the input database) of D(V/I)ExNPGP or G(V/I)ExNPGP, or octapeptides differing from these motifs at exactly one position, was calculated using the total number of octapeptides in the 131,941 amino acid sequences, and the relative frequencies of each of the 20 amino acids in this data set. It should be noted that these calculations assume that all of the sequences in the virus database are independent, which is not true (although heavily mitigated by the 70%-identity-threshold clustering step). Non-independence would tend to decrease the effective database size but, when and if false positives do occur, they may occur in multiple related sequences. It should be noted that, except at p12 and p15, the analyses of rare variants based on octapeptides that differ at exactly one position from D(V/I)ExNPGP (Fig. 4C) or G(V/I)ExNPGP (Fig. S4C) depend on completely non-overlapping sets of octapeptides, as those contributing to the former all contain D at p12 and those contributing to the latter all contain G at p12, yet the two analyses identify the same set of rare variants at p14, p16 and p19. The false positive rates (blue bars in Fig. 4C, S4C) represent the expected total number of octapeptides differing from D(V/I)ExNPGP or G(V/I)ExNPGP at the given specific position in randomized sequences. Thus, if all variations are non-functional (i.e. spurious), then the height of the blue bar is expected to approximate the sum of the heights of all of the red bars, excluding that for the canonical amino acid(s) (e.g. excluding G at p18). Our threshold of calling functional variants when the height of a single red bar is greater than three times the height of the blue bar is therefore very conservative.

Alphahelical propensity was predicted using AGADIR (25, 26) and NetSurfP3.0 (27). We used the 1900 representative 100 aa sequences divided into motif groups 1–6 besides the group of unclassifed sequences. One of the 100 aa sequences was removed due to the presence of unknown amino acids ("X"s). NetSurfP3.0 was accessed via the online server on 15 Mar 2025. AGADIR was downloaded on 17 Mar 2025 and run with parameters pH 7.2, temperature 310 K and ionic strength 120 mM. NetSurfP3.0 predicts three-category (helix, extended strand, coil) secondary structure of whole proteins, with a benchmark accuracy of ~0.60. Since the constrained environment of the exit tunnel prevents the formation of betasheets and other distal interactions, we also used AGADIR which assesses only local alphahelical propensity by modelling the thermodynamics of helix formation. Since distal interactions are irrelevant within the exit tunnel, we uniformly used 100 aa sequences flanking StopGo sites as input. The 12 terminal positions were trimmed from each end for plotting to mitigate edge-effects in the predictions. Although the sequence downstream of NPGP has not been synthesized when StopGo occurs, we included it to provide an illustrative base-line against which to compare the alphahelical propensity of the upstream sequence, noting that the StopGo octapeptide itself appears to provide a universal break in predicted alphahelical propensity.

To generate the codon usage pie charts, we used the 100 aa sequences flanking occurrences of (D/G/C/N)(V/I)ExNPGP restricting to sequences with defined host taxonomy, clustered these with BLASTCLUST with a 80% amino acid identity threshold, and selected a representative sequence from each cluster as described above, yielding 323 representative sequences of which 176 were from chordate hosts and 98 from arthropod hosts. The 80% instead of 50% identity threshold was used to provide more StopGo octapeptide examples (176 versus 123 and 98 versus 62, respectively), bearing in mind also that (in the absence of constraint) synonymous nucleotide substitutions generally accumulate more rapidly than amino acid substitutions. The *Homo sapiens* and *Drosophila melanogaster* codon usage statistics were downloaded from https://www.kazusa.or.jp/codon/ on 9 January 2025.

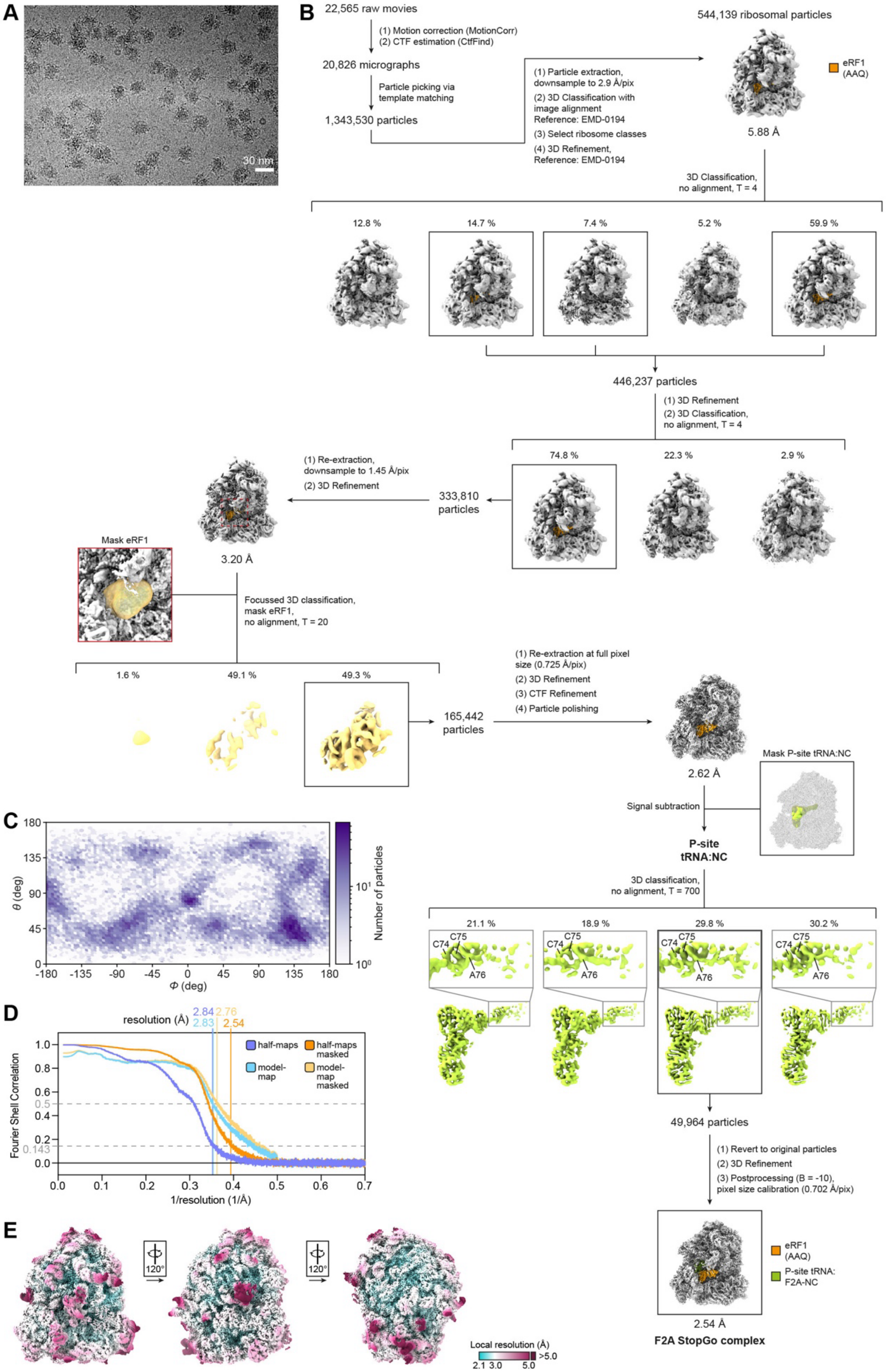
A
30 nm
B
22,565 raw movies
(1) Motion correction (MotionCorr)
(2) CTF estimation (CtfFind)
20,826 micrographs
Particle picking via template matching
1,343,530 particles
(1) Particle extraction, downsample to 2.9 Å/pix
(2) 3D Classification with image alignment Reference: EMD-0194
(3) Select ribosome classes
(4) 3D Refinement, Reference: EMD-0194
544,139 ribosomal particles
eRF1 (AAQ)
5.88 Å
3D Classification, no alignment, T = 4
12.8 %
14.7 %
7.4 %
5.2 %
59.9 %
446,237 particles
(1) 3D Refinement
(2) 3D Classification, no alignment, T = 4
74.8 %
22.3 %
2.9 %
(1) Re-extraction, downsample to 1.45 Å/pix
(2) 3D Refinement
333,810 particles
3.20 Å
Mask eRF1
Focussed 3D classification, mask eRF1, no alignment, T = 20
1.6 %
49.1 %
49.3 %
165,442 particles
(1) Re-extraction at full pixel size (0.725 Å/pix)
(2) 3D Refinement
(3) CTF Refinement
(4) Particle polishing
2.62 Å
Mask P-site tRNA:NC
Signal subtraction
P-site tRNA:NC
3D classification, no alignment, T = 700
21.1 %
18.9 %
29.8 %
30.2 %
C74
C75
A76
49,964 particles
(1) Revert to original particles
(2) 3D Refinement
(3) Postprocessing (B = -10), pixel size calibration (0.702 Å/pix)
eRF1 (AAQ)
P-site tRNA: F2A-NC
2.54 Å
F2A StopGo complex
C
θ (deg)
Φ (deg)
Number of particles
D
resolution (Å)
2.84
2.83
2.76
2.54
half-maps
half-maps masked
model-map
model-map masked
Fourier Shell Correlation
1/resolution (1/Å)
E
120°
Local resolution (Å)
2.1 3.0 5.0 >5.0

**Fig. S1. Cryo-EM data processing.** **(A)** Representative micrograph of the F2A StopGo cryoEM sample. The scale bar represents 30 nm. **(B)** Data processing pipeline. CryoEM data were first pre-processed (motion correction, CTF estimation, template-based autopicking), then the initial set of particles was subjected to 3D classification for identification of ribosomal particles. The particles were then further subjected to 3D classification without image alignment, followed by focussed classification for eRF1(-AAQ) (coloured orange). The resulting particles were then subjected to CTF refinement and polishing, before further focussed classification with signal subtraction for P-tRNA:NC. The Relion-reported resolutions of intermediate and final maps are indicated. **(C)** Euler angle distribution of the final set of particles, generated using the angdist script (https://github.com/Guillawme/angdist). **(D)** Map-map and map model Fourier shell correlation (FSC) curves. **(E)** Local resolution estimation. The map was generated using B-factor dependent sharpening and coloured according to local resolution (see colour code).

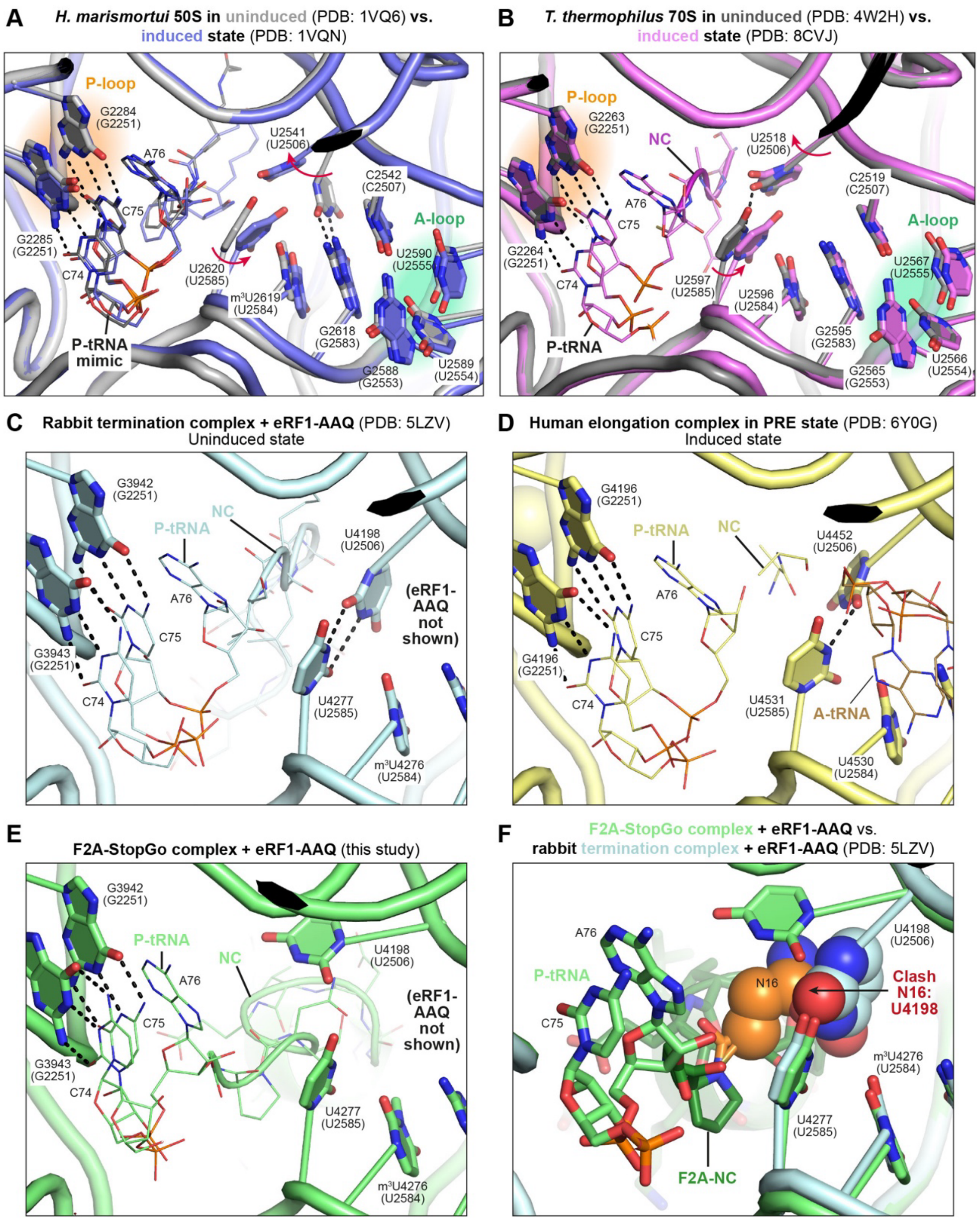


**Fig. S2. Conformation of PTC in the uninduced and induced states and comparison to the F2A-StopGo complex. (A), (B)** Superposition of the PTC structures of *H. marismortui* 50S in complex with tRNA analogues **(A)** or *T. thermophilus* 70S in complex with complete, acylated tRNAs **(B)**, modulating the ribosome PTC in either the induced or uninduced states. For clarity, only the tRNA (analogues) of the induced states are shown. **(C) – (E)** PTC structures of a rabbit termination complex with accommodated eRF1-AAQ (**C**, uninduced state), a human PRE complex after peptidyl transfer to the A-site (**D**, induced state), and the F2A-StopGo complex with eRF1-AAQ (**E**). **(F)** Superposition of structures of the F2A-StopGo complex and the canonical rabbit termination complex. The F2A-NC andU4198 are shown as spheres to highlight a clash between the RNA base and N16 of the NC. For all complexes, numbers in parentheses indicate the corresponding *E. coli* numbering as a reference.

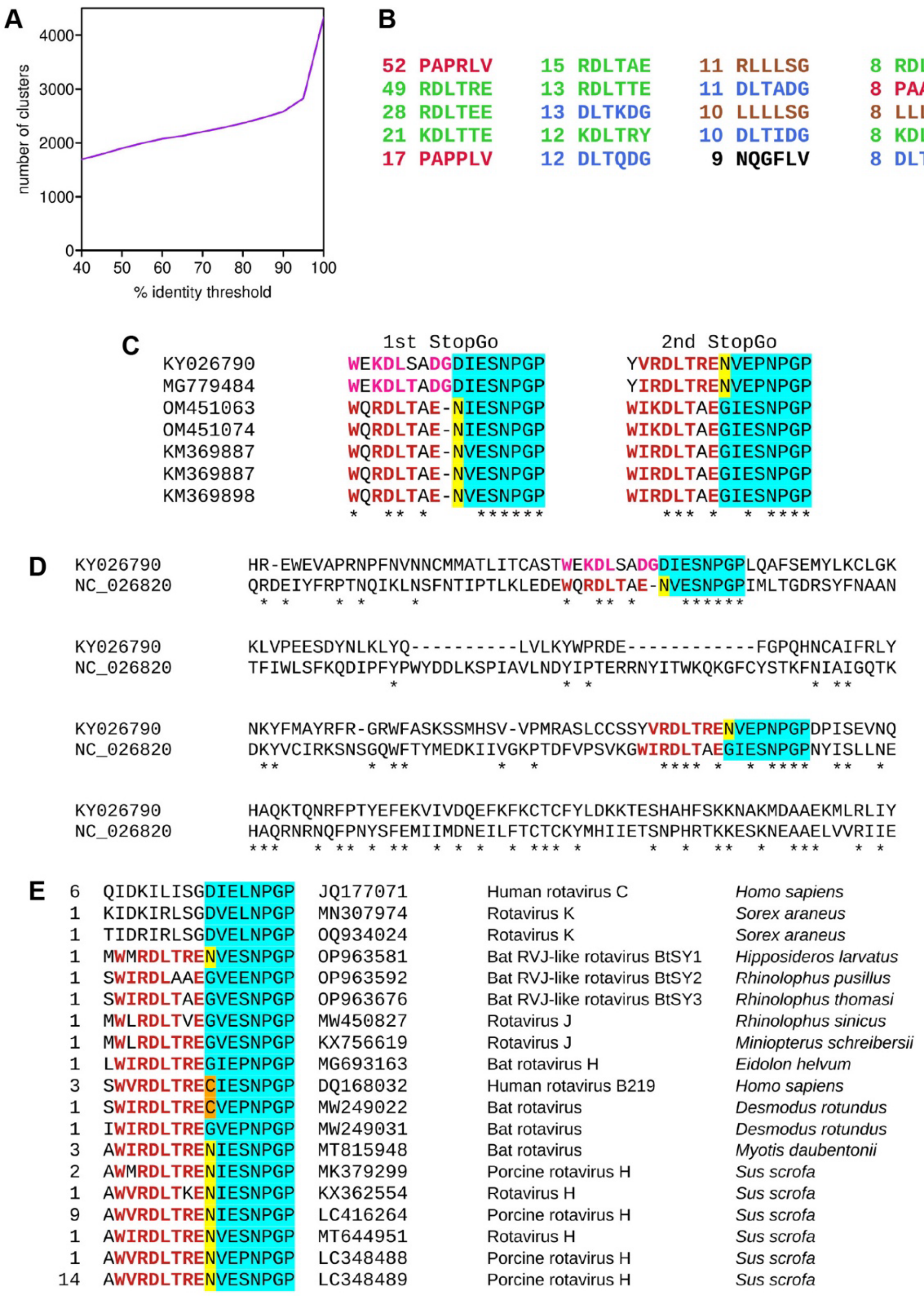


**Fig. S3. Identification of distinct motif classes upstream of the StopGo core octapeptide and examples supporting p12 = N and p12 = C as *bona fide* functional variants of StopGo. (A)** Sequence fragments comprising 100 aa surrounding (D/G/C/N)(V/I)ExNPGP octapeptides were clustered. The graph shows the number of BLASTCLUST clusters as a function of clustering identity threshold. **(B)** The 20 most abundant hexapeptides immediately upstream of the (D/G/C/N)(V/I)ExNPGP in the 1900 representative sequences. Numbers show the number of occurrences of each hexapeptide. Colours denote four groups of similar hexapeptides, while one singleton hexapeptide is shown in black. **(C)** C-terminal 240 columns of an alignment of the NSP1-2 amino acid sequences from rotavirus I NCBI accessions KY026790 and KM369887. Sequences were aligned with MUSCLE version 3.8.31 (28). These sequences are homologous (blastp, 2-sequence alignment, e-value = 5 x 10−61) but highly divergent (30% identity). Each sequence has two StopGos (core octapeptides highlighted in cyan). At each StopGo, one of the sequences has p12 = N (highlighted in yellow) whereas the other sequence has the more common p12 = D or G. Besides conservation, characteristic upstream motif 1 (bold red letters) or motif 4 (bold pink letters) amino acids further support these being functional StopGo sites. Note that full-length NSP1-2 is shorter than the 500 amino acid length threshold used in the main analysis, so all examples shown in this figure are in addition to the p12 = N and p12 = C octapeptides enumerated elsewhere. **(D)** Both sites are conserved in all rotavirus I NSP1-2 sequences currently available in NCBI, thus ruling out sequencing errors as an explanation for p12 = N. **(E)** Fifty other NCBI rotavirus accessions encoding (partial) homologues of rotavirus I NSP1-2 encode the second StopGo, whereas the first StopGo and adjacent domain appears to be unique to rotavirus I. The core StopGo octapeptide and upstream nine amino acids were extracted from each sequence. Each unique 17-mer is shown. The number of times each unique 17-mer appears among the 50 rotavirus sequences is listed at left, and one example accession number listed, with the virus name and source host species listed at right. Colour coding is as in panel A. Note that a number of other rotavirus sequences have p12 = N (yellow highlight) and two unique 17-mers (from four sequences in total) have p12 = C (orange highlight).

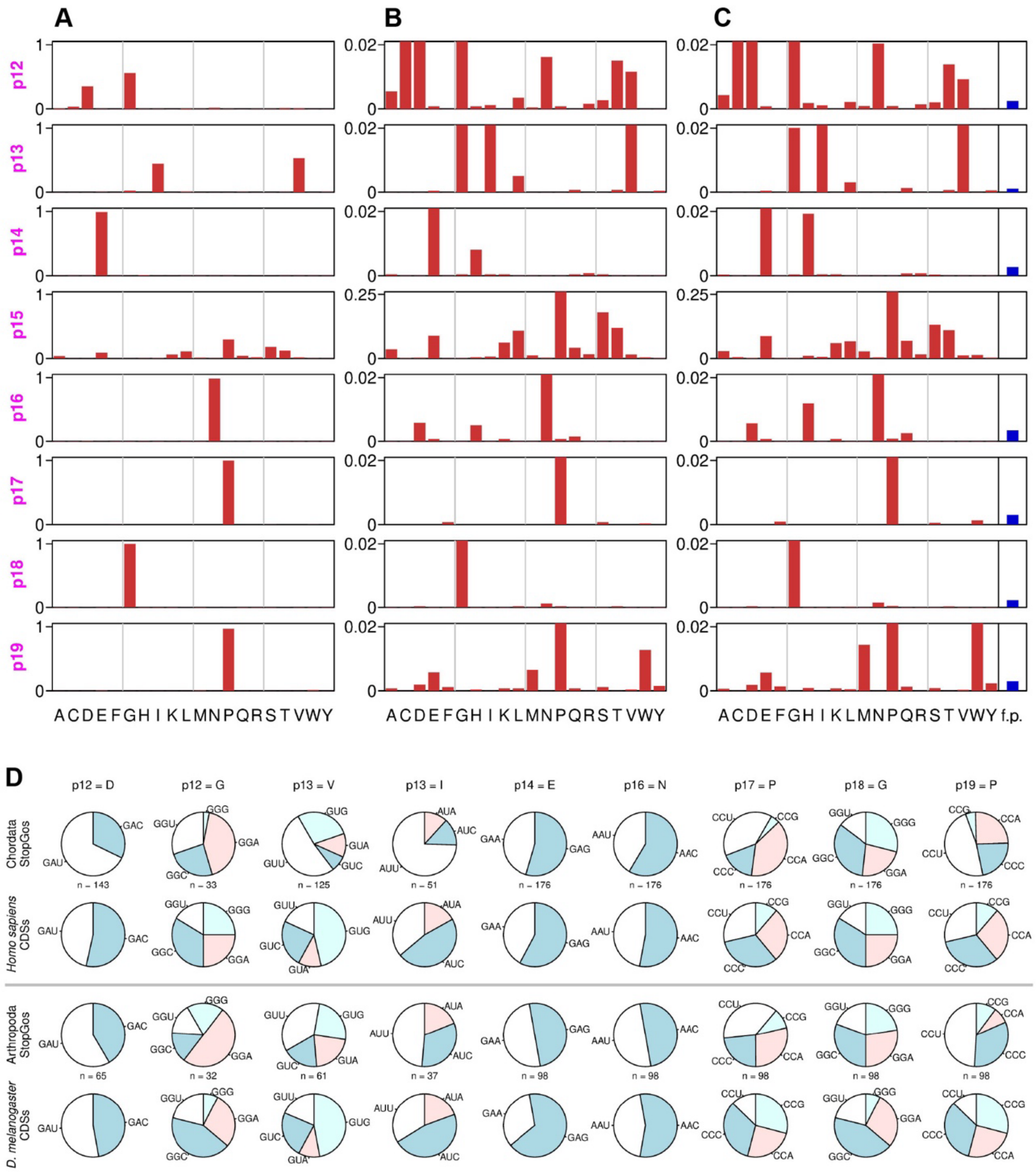


**Fig. S4. StopGo octapeptide variants and codon usage statistics. (A–C)** As for panels A–C in Figure 4, but using octapeptides differing from G(V/I)ExNPGP by at most one position (1207 exact matches; 1394 inexact matches). (**D**) Codon usage at the p12, p13, p14, p16, p17, p18 and p19 positions of 176 chordate and 98 arthropod virus StopGo octapeptides (rows 1 and 3, respectively). Note that *n* = 97 at Arthropoda p12, as one of the 98 octapeptides has p12 = N. Rows 2 and 4 show codon usage from representative host species *Homo sapiens* and *Drosophila melanogaster*.

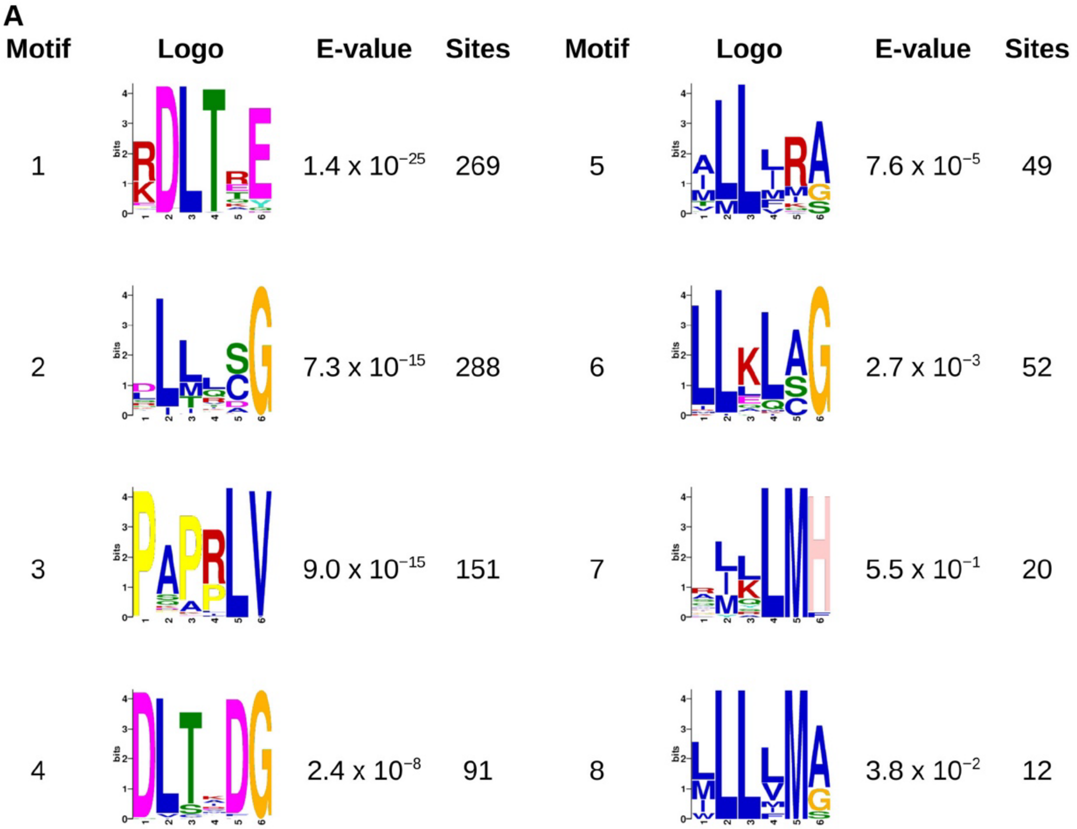


**B**

| | | | | |
|---|---|---|---|---|
| AB555545 | TLSGWIRDLTEEGVEPNPGPIVN | Omono River virus* | *Culex inatomii* | Arthropoda |
| ON812800 | GSCEWIRDLTEEGVEPNPGPGVS | Lonestar tick totivirus | *Hyalomma scupense* | Arthropoda |
| MN565680 | SKGGWIRDLTREGVEPNPGPVAG | Bremia lactucae associated fusagravirus 1 | *Bremia lactucae* | Oomycota |
| MZ209860 | TATGWVRDLTEEGVEPNPGPFGT | Sanya iflavirus 2* | *Sesamia inferens* | Arthropoda |
| KC843627 | TDKMWIRDLTQEGVEPNPGPHAE | Eel picornavirus 1 | *Anguilla anguilla* | Chordata |
| MW218667 | VEEGWVRDLTREGIEPNPGPVRV | Lanama virus* | *Chlorocebus pygerythrus* | Chordata |

**C**

| | | | | |
|---|---|---|---|---|
| MW218667 | AQGWVRDLTQDGDVESNPGPPQF | Lanama virus* | *Chlorocebus pygerythrus* | Chordata |
| OL956952 | NAAWVRDLTEDGDVEENPGPTQW | Duck egg reducing syndrome virus* | *Anas platyrhynchos* | Chordata |
| OL956952 | EHCWVRDLTMDGDVEENPGPWSP | Duck egg reducing syndrome virus* | *Anas platyrhynchos* | Chordata |
| ON746547 | DTDWVRDLTCDGDVEHNPGPISA | Yanbian Totiv tick virus 1 | *Haemaphysalis japonica* | Arthropoda |

**D**

| | | | | |
|---|---|---|---|---|
| AB035732 | FRSNYDLLKLCGDIESNPGPVTW | Bombyx mori cypovirus 1 | *Bombyx mori* | Arthropoda |
| OK422489 | QTLNFDLLKLAGDVESNPGPFFF | Foot-and-mouth disease virus O | *Bubalus bubalis* | Chordata |
| MG600092 | SFISRALLLLAGDVERNPGPCSF | Wenling bighead beaked sandfish picv.* | *Gonorynchus abbreviatus* | Chordata |
| MK645239 | EAARQMLLLLSGDVETNPGPVQS | Drosophila C virus | *Drosophila melanogaster* | Arthropoda |
| KX644936 | DPDLSSLLLLSGDVERNPGPCVI | Kunsagivirus B1 | *Eidolon helvum* | Chordata |

**E**

KYYYQPPAKRLVGIETNPGPWTP
VPPLFVPDPPLVGIETNPGPFNA
SYDQEVPAPRLVGIETNPGPDVP
NTYDDVPAPRLVGIEPNPGPPVL

**F**

| | | |
|---|---|---|
| M81861 | HYAGYFADLLIHDIETNPGPFMF | Encephalomyocarditis virus |
| X56019 | YHADYYKQRLIHDVEMNPGPVQS | Theilers murine encephalomyelitis virus |
| EF165067 | YHASYYKQRLQHDVETNPGPVQS | Saffold virus |
| KY432930 | YYKAFFDLKLQHDVETNPGPAQV | Cardiovirus F1 |
| MZ382838 | YYRQFCQNRLMHDVETNPGPVMS | Marmot cardiovirus |

**Fig. S5. Distinct motifs upstream of the StopGo core octapeptide. (A)** The top eight motifs identified by applying MEME to the six amino acids immediately upstream of the StopGo core octapeptide in the 1900 representative sequences. The table shows the sequence logo, E-value and number of sites reported by MEME. Due to their poor E-values, the last two motifs were discarded. **(B–D)** Examples of motif 1 (panel A; bold red letters), motif 4 (panel B; bold pink letters) and motifs 2 or 6 (panel C; bold green letters) in viruses from different host phyla. From left to right, columns show the NCBI accession number, StopGo octapeptide and flanking sequence, virus name, host species, and host phylum. Red asterisks indicate sequences that contain more than one StopGo site. **(E)** The four StopGos in sequence SRR12063586.macro.NODE_24_length_9085_cov_41.544326 from the rdrp_contigs.fa file of (4), derived from a pond water metatranscriptome (NCBI accession SRR12063586). Upstream amino acids characteristic of motif 3 are shown in bold orange letters. **(F)** Motifs 1–6 do not account for all StopGo cases (see unclassified [Un.] in Figure 5F,S6D). For example (and as noted previously by (29)), in cardioviruses there is a characteristic H at p11 – a feature which is not present in any of motifs 1–6. The upstream sequence in cardioviruses has similarities (bold mauve letters) with the discarded motif 7 from the MEME analysis (panel A).

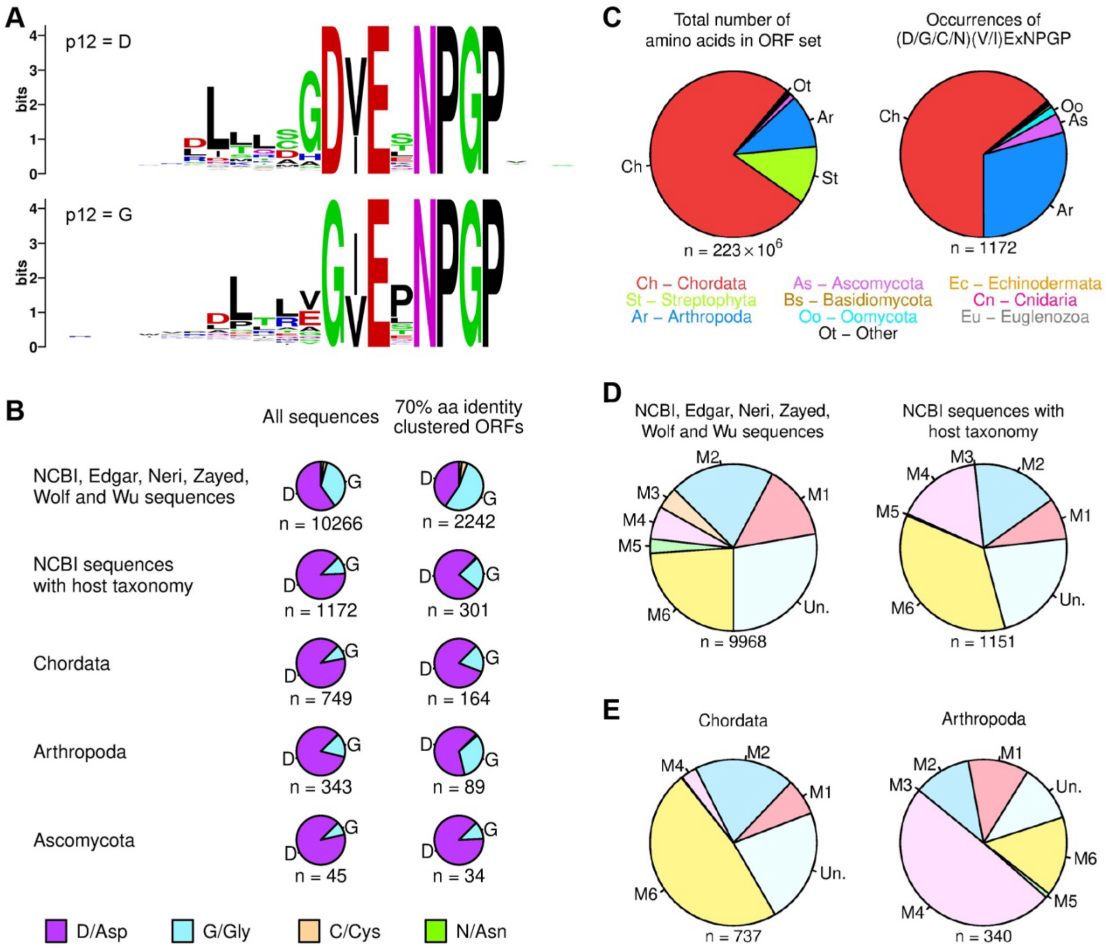


**Fig. S6. Analyses of StopGo position p12, and distribution of StopGo core octapeptides and the upstream hexapeptide motifs. (A)** Weblogos for sequences with p12 = D (top) or p12 = G (bottom). **(B)** Proportion of (D/G/C/N)(V/I)ExNPGP octapeptides with p12 = D, G, C or N, separated by sequence dataset or, where available, virus host taxon. **(C) – (E)** Host association of StopGo core octepaptide and upstream hexapeptide motifs as in Fig. 5E – G, but for the whole set of ORFs.

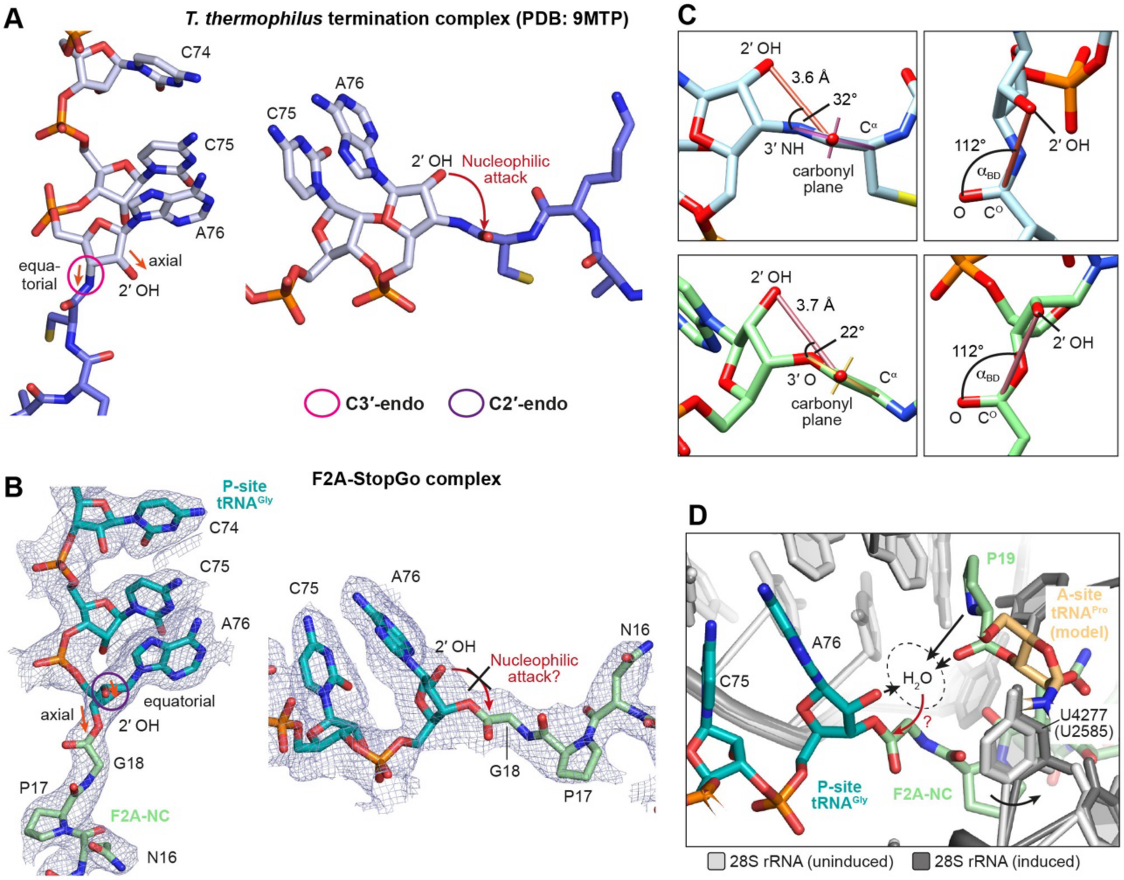


**Fig. S7. Putative mechanism for release of the StopGo NC. (A), (B)** Structures of the P-site tRNA CCA end and attached NC of a *T. thermophilus* termination complex (A; PDB ID: 9MTP) and the F2A-StopGo complex (B). The conformation of the A76 ribose 2′ and 3′ OH groups (axial versus equatorial) is indicated along with assignment as 3′ or 2′ endo sugar pucker. **(C)** Distances and angles of the A76 2′ OH group in relation to the ester carbonyl group in the *T. thermophilus* termination complex (top; PDB ID: 9MTP) or the F2A-StopGo complex (bottom). The carbonyl plane was defined using the $C^{\alpha}$,O and 3′ O/N atoms **(D)** Model of the PTC region in a model of the F2A-StopGo complex containing an acylated $tRNA^{Pro}$ in the A-site. The 28S rRNA is shown as seen in the experimental structure with eRF1-AAQ in the A-site, and after taking into account structural changes induced by tRNA binding to the A-loop (A-site tRNA and local rRNA changes obtained from PDB-model 6Y0G). Shift of the side chain of U4277 and the conformation imposed by the F2A-NC result in a large unoccupied volume (dashed) above the ester carbonyl group. The tRNAs and amino acyl moieties could contribute to coordination of a water molecule for nucleophilic attack on the ester bond. Numbers in parentheses indicate *E. coli* numbering.

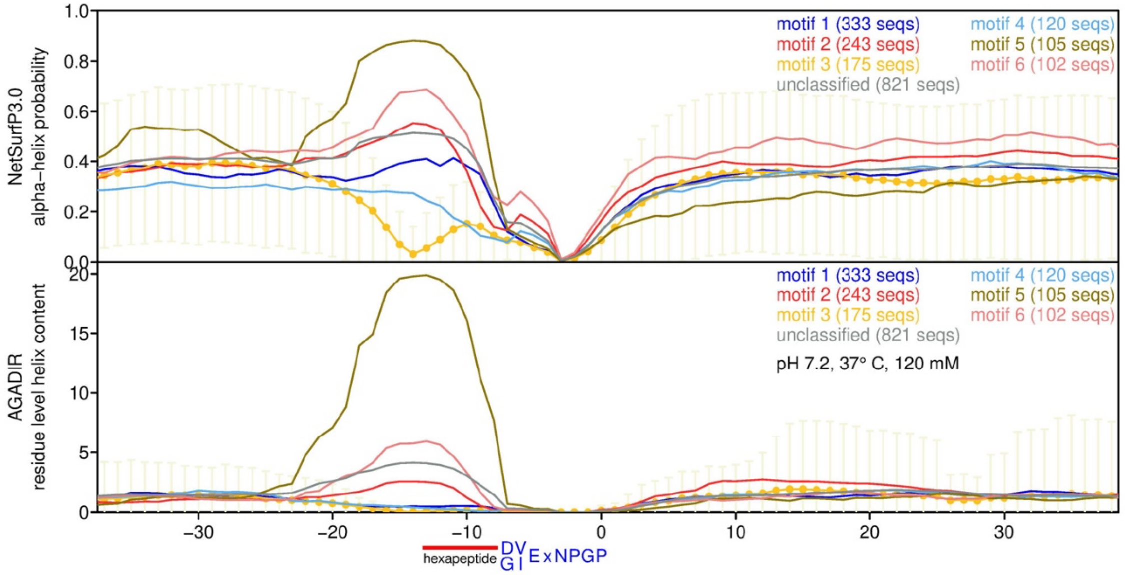


**Fig. S8. Analyses of StopGo alpha helical propensity.** Alpha helical propensity of StopGo-proximal sequences by motif group. NetSurfP3.0 alpha-helix probability score (upper) and AGADIR residue level helix content score (lower). Lines show mean values at each amino acid position across all sequences in a motif group. To avoid clutter, error bars (mean +/− s.d.) are shown for motif group 3 only.

**Table S1.** Cryo-EM data collection, refinement, and validation statistics for the F2A-StopGo complex

| | **F2A-StopGo complex**<br>(PDB 9RHU)<br>(EMDB EMD-53976) |
|---|---|
| **Data collection and processing** | |
| Microscope | FEI Titan Krios G1 |
| Voltage (keV) | 300 |
| Camera | Gatan K3 post-GIF direct electron detector + Gatan Quantum energy filter |
| Nominal magnification (x fold) | 105,000 |
| Pixel size at detector (nominal/calibrated) (Å/pixel) | 0.725 / 0.702 |
| Physical pixel dose rate ($e^{-}$/pixel/s) | 17.114 |
| Exposure time (s) | 1.2 |
| Total electron exposure ($e^{-}/Å^2$) | 40 |
| No. of frames collected during exposure | 89 |
| No. of fractions grouped | 40 |
| Dose per fraction ($e^{-}/Å^2$) | 1 |
| Defocus range (μm) | -2.2, -2.0, -1.8, -1.6, -1.2, -1.0 |
| Automation software | EPU v. 2.12.2 |
| Micrographs collected (no.) | 22,565 |
| Processing software | RELION 5.0.0 |
| Micrographs used (no.) | 20,826 |
| Total extracted particles (no.) | 544,139 |
| Final particles (no.) | 49,964 |
| Point-group or helical symmetry parameters | C1 |
| **Resolution estimates[1] (global, Å) (unmasked/masked)** | |
| Half-maps FSC = 0.143 | 2.84 / 2.54 |
| Model-map FSC = 0.5 | 2.83 / 2.76 |
| $d_{99}$ | 2.55 / 2.42 |
| $d_{model}$ | 2.60 / 2.60 |
| Resolution range[2] (local, Å) | 2.17 – 24.78 |
| Map sharpening *B* factor ($Å^2$) | -10 |
| **Model composition** | |
| Non-hydrogen atoms | 233,638 |
| Protein residues | 13,101 |
| RNA nucleotides | 5,976 |
| $Mg^{2+}$ ions | 367 |
| $K^{+}$ ions | 37 |
| $Zn^{2+}$ ions | 8 |
| $Fe_4S_4$ clusters | 2 |
| Spermidine | 2 |
| **Model refinement** | |
| Refinement package | phenix.real_space_refine_1.21.2-5149 |
| **Model-Map scores** | |
| CC (mask) | 0.87 |
| CC (volume) | 0.84 |
| **Average grouped B factors ($Å^2$)** | |
| Protein residues | 53.60 |
| RNA nucleotides | 62.96 |
| Ligands | 38.61 |
| **r.m.s.d. from ideal values** | |
| Bond lengths (Å) | 0.007 |

| | |
|---|---|
| Bond angles (°) | 1.021 |
| **Validation** | |
| MolProbity score | 1.21 |
| CaBLAM outliers (%) | 1.31 |
| Clashscore | 3.35 |
| Poor rotamers (%) | 0.10 |
| C-beta deviations (%) | 0.01 |
| **Ramachandran plot** | |
| Favored (%) | 97.61 |
| Allowed (%) | 2.35 |
| Outliers (%) | 0.04 |
| **Ramachandran plot *Z*-score, (r.m.s.d.)** | |
| whole | -0.88 (0.07) |
| helix | -0.54 (0.07) |
| sheet | -0.20 (0.11) |
| loop | -0.62 (0.07) |

[1] Calculated using comprehensive validation tool within the Phenix suite (mask based on model)
[2] Calculated using Relion's local resolution estimation
